# High-throughput DNA extraction and cost-effective miniaturized metagenome and amplicon library preparation of soil samples for DNA sequencing

**DOI:** 10.1101/2023.09.04.556179

**Authors:** Thomas BN Jensen, Sebastian M Dall, Simon Knutsson, Søren M Karst, Mads Albertsen

## Abstract

Reductions in sequencing costs have enabled widespread use of shotgun metagenomics and amplicon sequencing, which have drastically improved our understanding of the microbial world. However, large sequencing projects are now hampered by the cost of library preparation and low sample throughput. Here, we benchmarked three high-throughput DNA extraction methods: ZymoBIOMICS™ 96 MagBead DNA Kit, MP Biomedicals^TM^ FastDNA^TM^-96 Soil Microbe DNA Kit, and DNeasy® 96 PowerSoil® Pro QIAcube® HT Kit. The DNA extractions were evaluated based on length, quality, quantity, and the observed microbial community across five diverse soil types. DNA extraction of all soil types was successful for all kits, however DNeasy® 96 PowerSoil® Pro QIAcube® HT Kit excelled across all performance parameters. We further used the nanoliter dispensing robot I.DOT One to miniaturize Illumina amplicon and metagenomic library preparation volumes by a factor of 5 and 10, respectively, with no significant impact on the observed microbial communities. With these protocols, DNA extraction, metagenomic library preparation, or amplicon library preparation for one 96-well plate are approx. 3, 5, and 6 hours, respectively. Furthermore, the miniaturization of amplicon and metagenome library preparation reduces the chemical and plastic costs from 5.0 to 3.6 and 59 to 7.3 USD pr. sample.

## Introduction

The drastic reduction in sequencing costs has enabled more researchers to utilize next-generation sequencing in their field of study (1). In the field of microbial ecology especially, the reduced costs have enabled an increase in the scope of the projects, as thousands of samples are required to understand the diversity of microbial habitats (2–5). The ongoing reductions in sequencing costs means that large sequencing projects are now cost-limited by the cost associated with hands-on time and sample preparation. However, new automated or semi-automated workflows utilizing liquid handlers and drop dispensing technology seem promising regarding the reduction of both labor time and reaction volumes - ultimately reducing costs (1,6–8).

Soil samples are especially problematic for high-throughput (HT) DNA extraction workflows due to the diversity of soil properties (1,9,10). Furthermore, the majority of the proposed protocols are not easy to convert to a HT format due to steps that are either difficult to automate, time-consuming, or include hazardous substances. Although several commercial soil-specific HT DNA extraction kits are available (table 1), these have not been independently tested on a diverse range of soil types.

**Table 1.** Commonly used and commercially available kits for DNA extraction from soil. Extraction kits in **bold** were compared. High-throughput equipment refers to available automated solutions for the HT solution. LT: Low-throughput, HT: High- throughput.

| Manufacturer | Low-throughput Kit | High-throughput Kit | High-throughput Equipment | Reference |
| --- | --- | --- | --- | --- |
| QIAGEN<br>(previously MoBio) | **DNeasy® PowerSoil® Kit\***<br>(PowerSoil LT) | DNeasy® PowerSoil® HTP 96 Kit |  | LT: (10–21)<br>HT: (22) |
|  |  | MagAttract® PowerSoil® Pro DNA Kit | KingFisher® Flex/Duo<br>epMotion® 5075 TMX | HT: (5,22,23) |
|  | DNeasy® PowerSoil® Pro Kit | **DNeasy® 96 PowerSoil® Pro QIAcube® HT Kit (Powersoil Pro HT)** | QIAcube® HT System | HT: (24–26) |
| MP Biomedicals | **FastDNA™ SPIN kit for Soil**<br>(FastSpin LT) | **FastDNA™-96 Soil Microbe DNA extraction Kit**<br>(FastSpin HT) |  | LT: (10,19,20,27,28) |
| Macherey-Nagel | NucleoSpin™ Soil Kit | NucleoSpin™ 96 Soil Kit |  | LT: (21,29–32)<br>HT: (33–36) |
| Zymo Research | ZymoBIOMICS® DNA Micro/Mini/Midi Kit | ZymoBIOMICS® 96 DNA Kit |  | LT: (16,17)<br>HT: (37) |
|  |  | **ZymoBIOMICS® 96 MagBead DNA Kit**<br>(ZymoMagbead HT) | Hamilton Microlab® STAR | HT: (22,23) |
|  | ZymoBIOMICS®<br>Quick-DNA Fecal/Soil<br>Microbe<br>Micro/Mini/Midi Kit | Quick-DNA Fecal/Soil Microbe<br>96 Kit |  |  |
|  |  | Quick-DNA Fecal/Soil Microbe<br>96 Magbead Kit | Hamilton Microlab® STAR | HT: (38) |
| Omega Bio-Tek | E.Z.N.A.® Soil DNA Kit | Mag-Bind® Environmental DNA<br>96 Kit | Hamilton Microlab® STAR<br>Hamilton Microlab® NIMBUS<br>KingFisher™<br>BioSprint®<br>MagMAX® 96 | LT: (21,39–42) |

\*QIAGEN has replaced this kit with DNeasy PowerSoil Pro. C2 and C3 have been replaced by CD2 which is a combination of the two buffers.

With the reduction in sequencing costs, library preparation has become a significant proportion of total project costs. One of the first kits for cost-effective next-generation sequencing of low input material, which allowed for multiplexing of a large number of samples, was the Nextera XT DNA library preparation kit (43–45). The Nextera XT library preparation protocols for small genomes, PCR amplicons, plasmids, or cDNA have undergone several transformations since the first release of the Nextera XT protocol in 2012. The first Nextera XT protocols were easy to use, however the expensive reagents were limiting for large sequencing projects (46). To reduce the library preparation costs, earlier work focused on diluting the expensive reagents or replacing them with cheaper alternatives (47), however in 2017 the Nextera Flex (renamed to Illumina DNA prep) protocol was introduced, which utilizes bead-linked transposases, rendering the previous cost-effective protocols less useful. Previous work has shown it is also possible to dilute the reagents in the Nextera Flex kit (46), however another strategy for reducing the overall costs without tampering with the reagents is by miniaturization.

Here we present and benchmark a complete HT workflow from DNA extraction to miniaturized Illumina amplicon or metagenome library (Fig 1).

**Fig 1.**
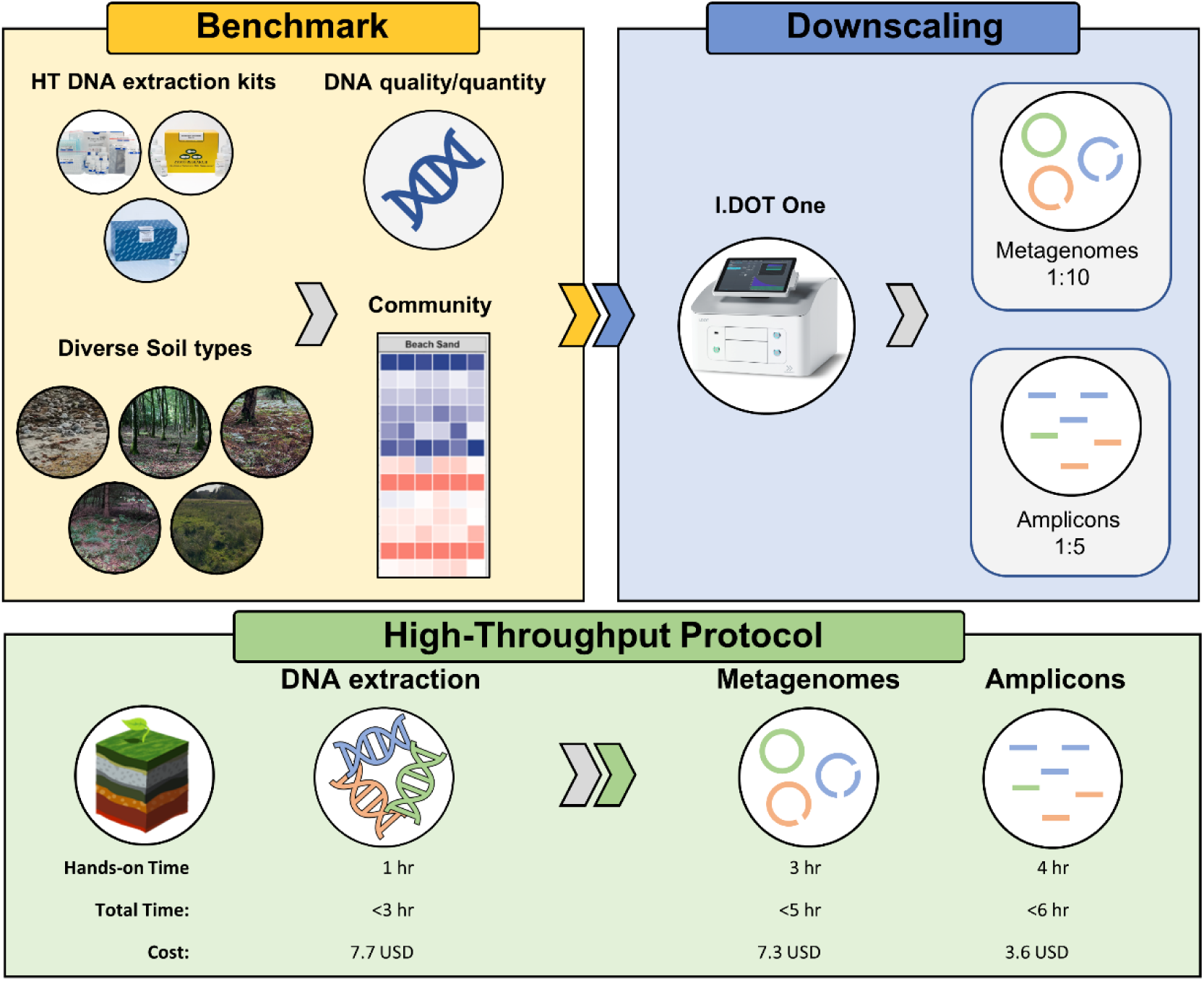
Experimental design. Top: Experimental design. Five different soil types were used to benchmark three different HT DNA extraction methods. DNA extractions were evaluated by DNA quality, quantity, length, and observed community profile. The I.DOT One was subsequently used to miniaturize metagenomes and amplicons. Bottom: Hands-on time, total time and cost associated with each step from DNA extraction to metagenomes or amplicons. The time reflects the processing time of a full 96- well plate, whereas costs are calculated per sample. For both metagenomes and amplicons, two 96-well plates can be processed concurrently with only minor added total time.

## Results

### 1. DNA Extraction Benchmark

The three HT DNA extraction kits were benchmarked on five different soil types (S1 Table). PowerSoil Pro HT and ZymoMagbead HT both come without a lysing matrix, whereas FastSpin HT, FastSpin LT, and PowerSoil LT do. To better evaluate the DNA extraction chemistry of the HT kits, it was decided to use the lysing matrix E from FastSpin LT. DNA extraction with PowerSoil LT was performed with its native lysing matrix. Samples were bead-beaten for six minutes total at 1800 RPM with the lysing matrix E. All samples were prepared according to the manufacturer’s protocol.

Generally, the kits were able to extract DNA from all soil types; however low amounts were extracted from Beach Sand, which was likely due to low biomass relative to the other soil types (S2 Table). PowerSoil Pro HT extracted more DNA than both FastSpin HT and ZymoMagbead HT (p<0.001, ANOVA on ranks, n=45). DNA yield of PowerSoil Pro HT and PowerSoil LT were comparable (p=0.08, ANOVA on ranks, n=30). The FastSpin LT DNA yields could not be determined due to unreliable Qubit measurements likely caused by a high concentration of residual humic substances after DNA extraction (Fig1 in S1 file).

Generally, the 260/280 ratios were around ∼1.8, which is considered pure for DNA. The FastSpin HT had high 260/280 ratios for all soil types, which could indicate contamination with RNA. The 260/230 ratio varied greatly between kits. Low values were measured for FastSpin LT, FastSpin HT, and ZymoMagbead HT indicating the inability to remove sample or kit contaminants absorbing at 230 nm (S2 Table). The Powersoil Pro HT and Powersoil LT had 260/230 ratios closest to that of pure nucleic acids, which is 2.0-2.2. The 260/230 ratio of PowerSoil Pro HT was not significantly different from its LT counterpart (Mann-Whitney U, p=0.35), but a higher 260/280 ratio was observed (Mann-Whitney U, p<0.001).

DNA shearing among the HT kits was highest for PowerSoil Pro HT with a mean peak DNA fragment length of 7.3 kb (n=15, sd = 0.7 kb) compared to an average length of 13.2 kb and 16.0 kb for FastSpin HT (n=15, sd=3.9 kb) and ZymoMagbead HT (n=15, sd=5.4 kb), respectively (S1 Fig). DNA shearing was primarily driven by the extraction kit (81.7 % variance, p<0.001, ANOVA on ranks n=75) but also the interaction of the extraction kit and soil type (10.8 % variance, p<0.001, ANOVA on ranks n=75). The mean peak fragment length of the FastSpin LT kit, 5.6 kb (n=15, sd=0.9 kb), was even lower than for PowerSoil Pro HT. PowerSoil LT mean peak fragment length, 8.6 (n=15, sd=1.4 kb), was significantly larger than that of the PowerSoil Pro HT kit (Mann-Whitney U, p<0.001).

Miniaturized Illumina amplicon libraries were sequenced and processed through standard bioinformatic pipelines (see methods). Two out of three organic soil amplicon libraries failed for FastSpin LT and ZymoMagbead HT. Both kits resulted in low 260/230 ratios suggesting the presence of contaminants. Amplicon libraries were successfully sequenced for all soil types for both FastSpin HT and Powersoil Pro HT; however, with both methods one library from the Sand samples was removed as an outlier evaluated by principal component analysis (PCA) - most likely due to contamination of the samples.

Based on ANOVA, the Shannon diversity index was significantly different for both soil type (83.2 % variance explained, p<0.001), DNA extraction kit (7.1 % variance explained, p<0.001), and the interaction of soil type and kit (6.7 % variance explained, p<0.001). The difference between PowerSoil Pro HT and FastSpin HT was 0.04 (Tukey’s HSD, p=0.03), between FastSpin HT and ZymoMagbead was 0.05 (Tukey’s HSD, p=0.02), and between PowerSoil Pro HT and ZymoMagbead HT was 0.09 (Tukey’s HSD, p<0.001). Similarly, Bray-Curtis dissimilarity was significantly different for both soil type (61.5 % variance explained, p<0.001), the interaction of soil type and kit (23.2 % variance explained, p<0.001), and for the extraction kit (6.1 % variance explained, p<0.001).

The microbial community profiles were similar across all kits (Fig 2A). PCA revealed the communities clustered according to soil type, not DNA extraction kits. Based on a PERMANOVA 2.8 % variance was explained by DNA extraction kit (p<0.001) and 90.6 % by soil type (p<0.001) (Fig 2B). When stratifying for soil type PCA revealed samples clustered by DNA extraction kit (Fig 2C, S2 Fig).

**Fig 2.**
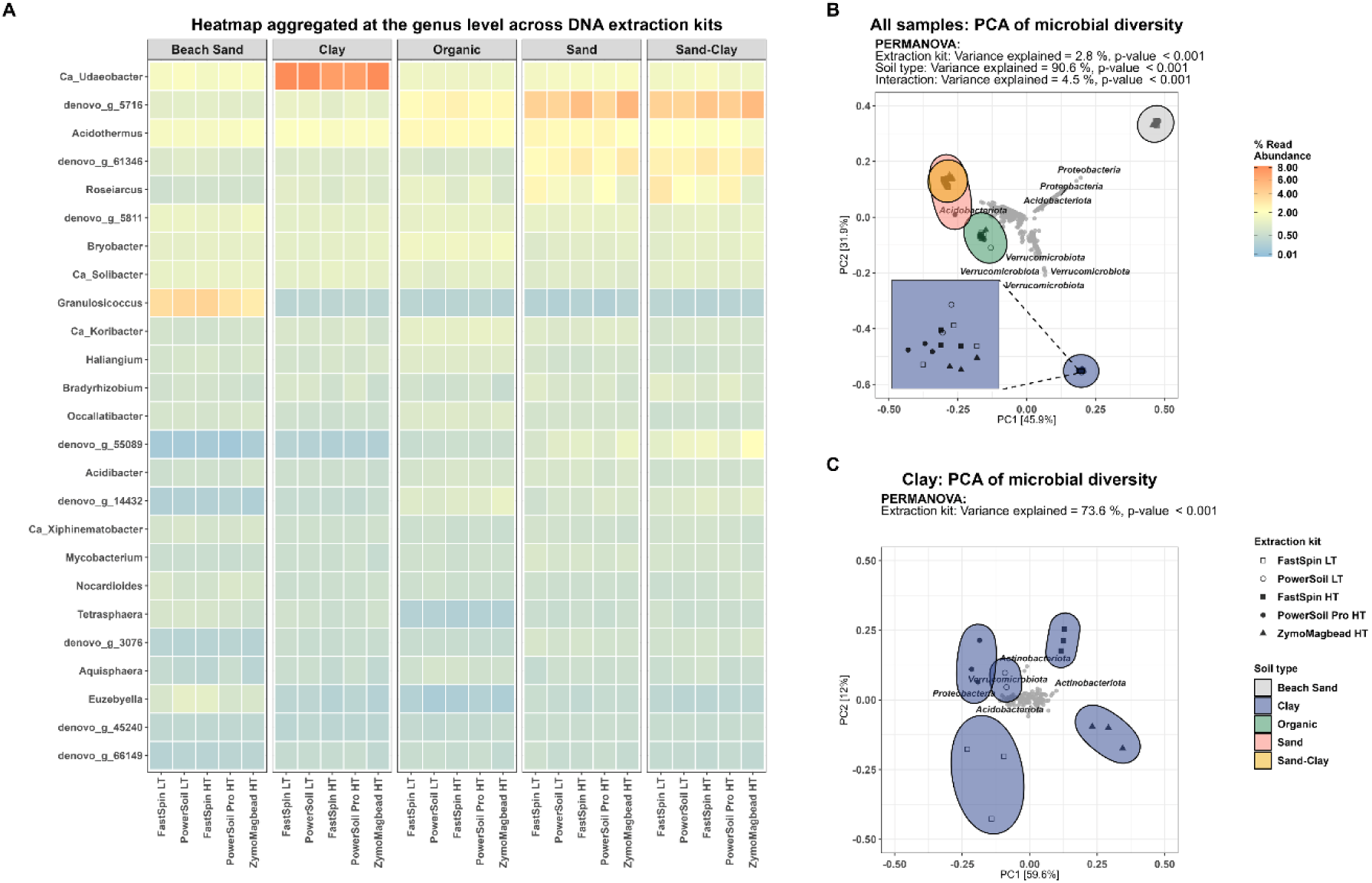
Community characteristics for DNA extraction kits. (A) Heatmap of community profile at phylum level across DNA extraction kits, faceted by soil type. (B) PCA on Hellinger transformed relative abundance. (C) PCA on Hellinger transformed relative abundance for the Clay samples only. For all plots: ASVs not exceeding 0.1 % relative abundance in at least one sample were filtered out.

Based on the DNA extraction characteristics, the high amplicon library success rate for all soil types, and the consistent community profile the PowerSoil Pro HT DNA extraction kit was selected for further optimization. Specifically, the effect of bead beating time and intensity on the observed community structure and DNA quantity and length was investigated. Both bead-beating time and intensity affected the DNA yield and observed microbial community, however, little difference was observed between six minutes of bead-beating at 1600 or 1800 RPM. Increasing the bead-beating intensity to 1800 RPM did however increase fragmentation, therefore a bead-beating of a total of six minutes at 1600 RPM was chosen (S2 File). Reducing the input amount from 125 mg to 50 mg had no effect on the observed microbial community (S2 File). The PowerSoil Pro HT kit can furthermore be semi-automated with the QIAcube HT system to reduce the hands-on time. For our projects, the DNA extraction costs per sample based on chemicals and disposables for PowerSoil Pro HT was 7.7 USD.

### 2. Miniaturized Illumina Amplicon Library Protocol

Amplicon libraries were successfully prepared for all soil types (S3 Table). Shannon diversity index was not significantly affected by the library volume (0.3 % variance explained, p=0.33, ANOVA, n=30) when blocking the contribution from soil type (p<0.001) and the interaction between soil type and library volume (p=0.04). The data violate the assumption of normal distribution but not the assumption of heteroscedasticity for both grouping factors. ANOVA based on ranks did yield very similar results but changed the p-value of the interaction to p=0.14.

Mean Bray-Curtis dissimilarity between replicates was not significantly affected by library volume (p=0.86, ANOVA on ranks, n=30). Furthermore, the Bray-Curtis dissimilarity between protocols was similar to the dissimilarity between replicates with the exception being Beach Sand (p=0.002, Mann-Whitney U=15, Bonferroni correction for multiple testing).

Community profiles at the genus level were similar between standard and miniaturized amplicon libraries (Fig 3A). When conducting a differential abundance analysis of all ASVs between the standard and miniaturized protocol a total of 19 ASVs were found to be differentially abundant (Fig 3B). In total, 11 of these (8 from Beach Sand and 3 from Organic) were completely absent across all replicates in the miniaturized protocol, likely showcasing the limitations of miniaturization for low biomass and/or highly diverse samples. No differential abundant ASVs were detected in Sand, Sand-Clay, and Clay (S3 Fig).

**Fig 3.**
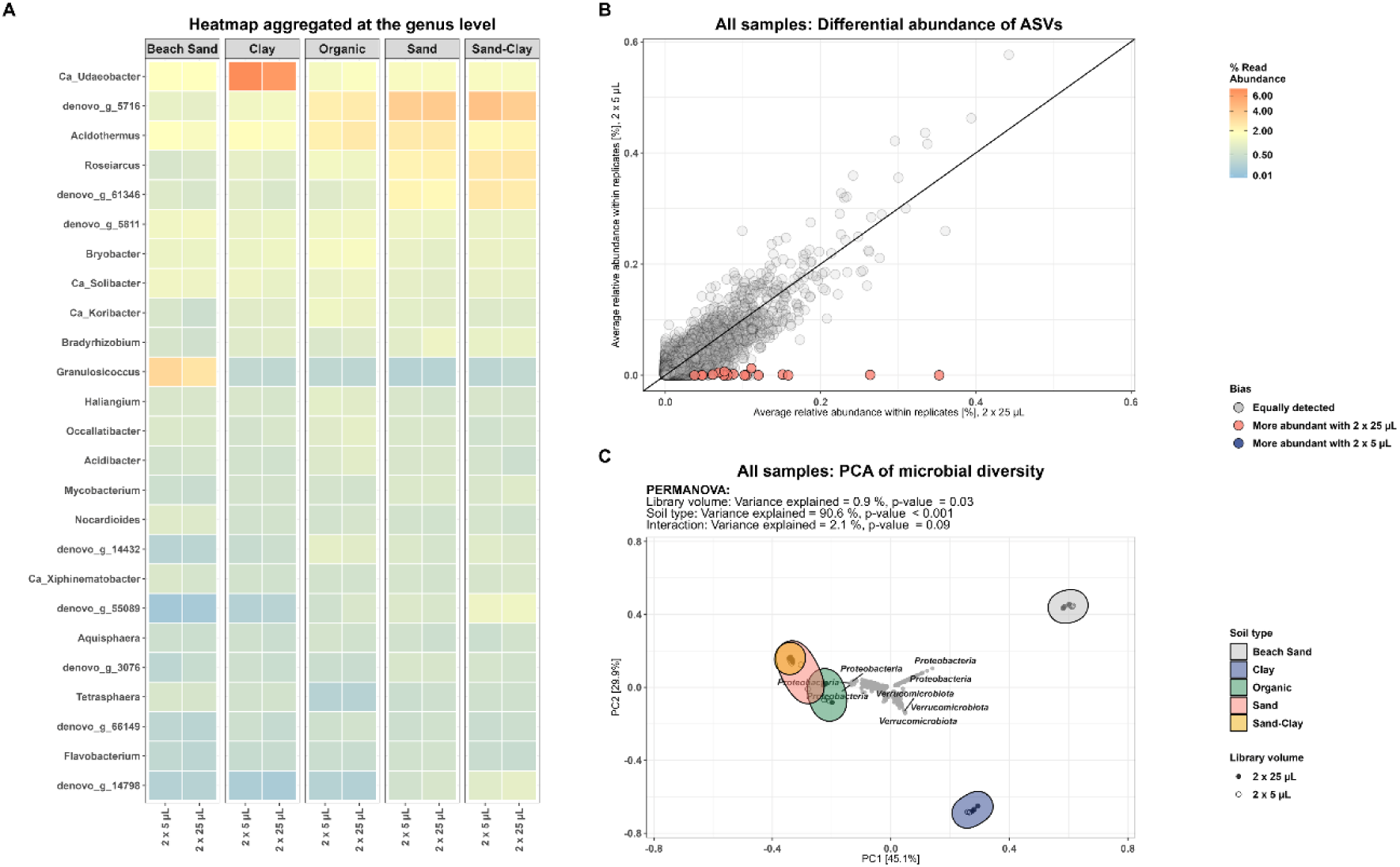
Comparison of community characteristics between standard and miniaturized amplicons. (A) Heatmap of community profile at genus level across reaction volume, faceted by soil type. (B) Differential relative abundance plot of ASVs. (C) Hellinger transformed relative abundance, PCA. ASVs not exceeding 0.1 % relative abundance in at least one sample were filtered out.

Using only ASVs with a relative abundance above 0.1 % the variance explained by the different library preparation protocols amounts to 0.9 % of the total variance observed (PERMANOVA, p=0.03) (Fig 3C). PERMANOVA showed no significance for library volume when performed per soil type (S4 Fig B-F).

The miniaturized protocol effectively reduced our chemical and plastic cost for amplicon library preparation from 4.9 USD to 3.6 USD.

### 3. Miniaturized Illumina DNA Prep Protocol

Illumina DNA prep libraries were successfully prepared and sequenced for all soil types with both the standard and the miniaturized protocol (S4 Table) except one library for the Clay soil using the miniaturized protocol.

All reads identified as 16S fragments were aggregated to the genus level (see methods). Based on ANOVA on ranks, Shannon diversity index was not significant for the library volume (0 % variance explained, p=1). Soil type explained 80.2 % variance (p<0.001). From the ANOVA analysis based on ranks, Bray-Curtis dissimilarity between replicates was affected by the library volume (6.8 % variance explained, p=0.02), being slightly lower in the miniaturized protocol (TukeyHSD, p=0.02). Soil type again accounted for the largest proportion of the variance (65 % variance explained, p<0.001). Bray-Curtis dissimilarity for replicates within protocols was not different to the dissimilarity of replicates between protocols for any soil type (S4 Table).

Comparison of the relative abundance at the genus level revealed very similar profiles between the standard and miniaturized protocol, regardless of soil type (Fig 4A). None of the identified genera were found to be differential abundant between the protocols (Fig 4B, S5 Fig). Using only genera with a relative abundance above 0.1 % the variance explained by the different library preparation protocols amounts to 0.3 % of the total variance observed (PERMANOVA, p=0.21) (Fig 4C). When performing PERMANOVA per soil type no significance for library volume was observed (S6 Fig B-F).

**Fig 4.**
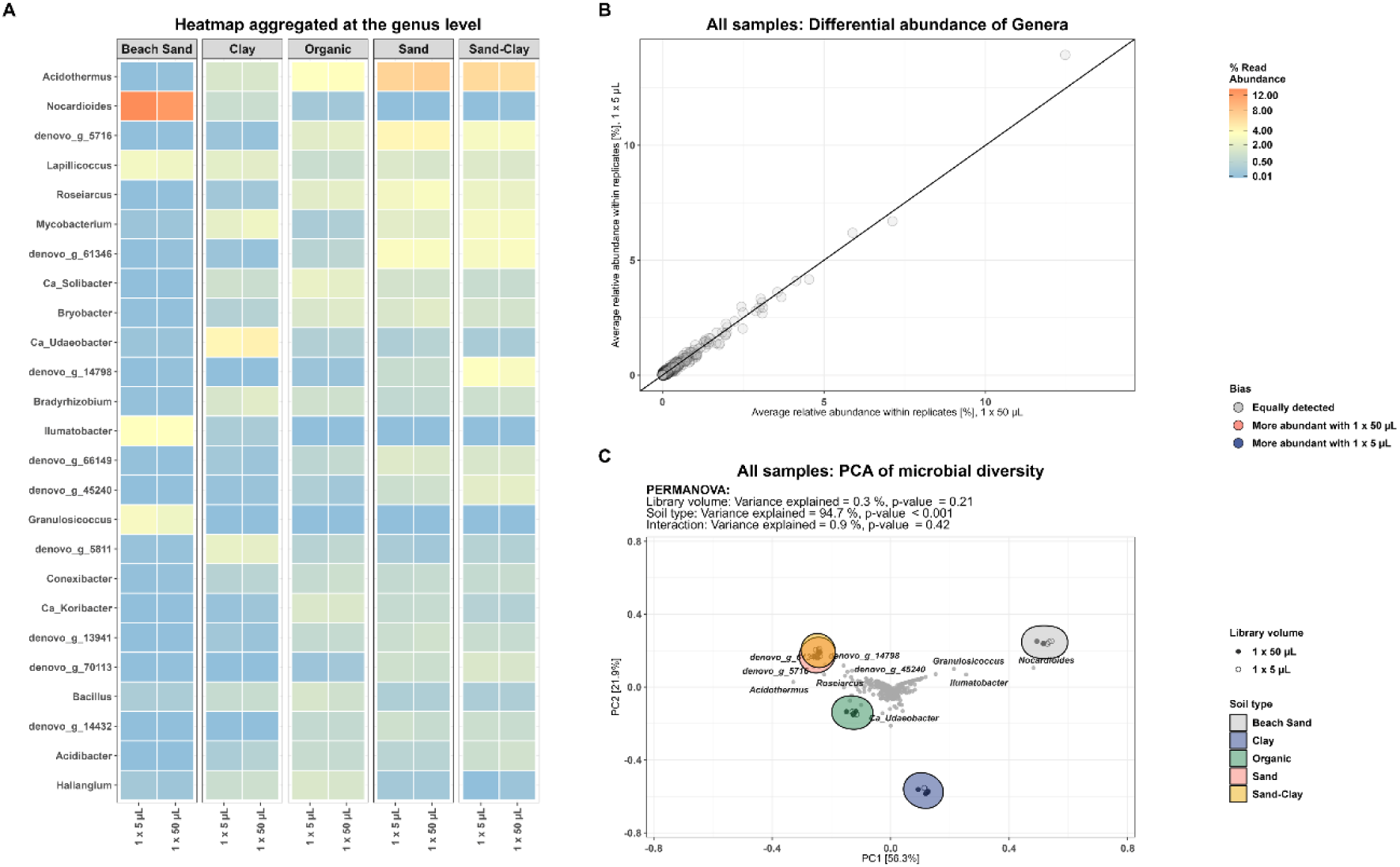
(A) Heatmap of community profile at genus level across reaction volume, faceted by soil type. (B) Differential abundance plot of all genera (C) Hellinger transformed relative abundance, PCA. Genera which did not exceed 0.1 % relative abundance in at least one sample were filtered out.

As part of the optimization step of the protocol, different ratios of Sample Purification Beads (SPB/IPB) were tested for optimal fragment size distribution, as a shift towards shorter fragments was observed with the miniaturized protocol (see S3 File). Miniaturizing the Illumina DNA library prep with a factor of 10 reduced our chemical and plastic cost for preparation of metagenomes from 52.2 USD to 7.3 USD.

## Discussion

Microbial communities are often entangled with questions which require large sample sizes to answer. Hence, to facilitate this it was paramount that sample preparation was converted to a HT setting and sample preparation costs are reduced to enable more large scope projects (2). For soil samples, the sheer amount of diversity poses a problem for DNA extraction kits, as the kit often needs to be optimized for each sample type, which is infeasible in a HT setting. In this benchmark, PowerSoil Pro HT slightly outperformed other HT kits for all soil types based on DNA quality, specifically the 260/230 ratio. Another advantage of the PowerSoil Pro HT was the semi-automated QIAcube extraction protocol, which significantly reduces hands-on time and the bias associated with manual liquid handling. Amplicon libraries for all soil types were successfully sequenced with DNA from the PowerSoil Pro HT (n=15) protocol and FastSpin HT (n=15). PowerSoil Pro HT did on average fragment the DNA more than other protocols, which can be a disadvantage for long-read DNA sequencing technologies, such as Oxford Nanopore Technology and PacBio. Though the peak fragment size was longer than the golden threshold of 7 kb (48) for all soil types most of the DNA was below this threshold.

The microbial community could successfully be analyzed with miniaturized reaction volumes for both amplicons and metagenomes. Metagenomes could be prepared in a 1:10 reaction volume, whereas amplicon reaction volumes could be miniaturized with a factor of five. A downside to the miniaturized reaction volumes was the entry cost of the nano-liter drop dispensing platforms, ranging from 100.000 to 300.000 USD, as well as expensive servicing fees and highly priced plastic consumables. However, in large projects, the entry cost was small compared to the reduction in library preparation cost and hands-on time. In our case, the cost savings of miniaturizing the metagenomes library protocol exceeded the price of the I.DOT One after ∼2000 samples.

Several steps were automated with the QIAcube HT and I.DOT One to reduce the hands-on time. The protocol could be further improved by automating some of the liquid handling steps, especially the clean-up steps in the HT miniaturized metagenomes protocol.

The cost of sequencing will depend on the question to be answered as some projects will require higher sequencing depth than others. If metagenomic samples are sequenced to a median depth of 5 Gbp with the Illumina NovaSeq 6000, it corresponds to approximately 19 USD in sequencing chemical costs per sample. For amplicons, multiplexing the maximum amount of samples (384 with current Illumina barcodes) for sequencing on the Illumina NovaSeq 6000 platform would give excessive depth even for the lowest throughput flow cell available, SP, amounting to ∼9-12 USD per sample in sequencing costs and a theoretical median depth of 1.69 to 2.08 million ∼250-450 bp amplicons. Amplicon sequencing costs could be reduced tremendously by designing additional barcodes for further multiplexing (46,49). Acquiring 10.000 unique (100 x 100) indexes of 500 picomole each, and sequencing on the SP flow cell would yield 65-80 thousand amplicons per sample, and amount to 48 cents for 250 bp amplicons and 62 cents for 450 bp amplicons.

## Conclusion

The DNeasy® 96 PowerSoil® Pro QIAcube® HT Kit chemistry outperformed both FastDNA™-96 Soil Microbe DNA extraction Kit and ZymoBIOMICS® 96 MagBead DNA Kit based on DNA purity and yield when using the lysing matrix E, though at the cost of more fragmented DNA. Metagenomes and amplicons could successfully be miniaturized without affecting the observed microbial community for five different soil types, effectively reducing the chemical and plastic costs for library preparation to 7.3 USD and 3.6 USD, respectively.

## Materials and methods

### Soil types

Five different soil types were included. All samples consist of five topsoil subsamples (0-20 cm) taken within five areas of ∼80 m2 (5 m radius), which were mixed before pouring into a 100 mL sample container. pH was measured for each soil type by mixing 10 g of soil with 30 mL of deionized water (50). After settling, pH was measured with [SI Analytics Lab 855]. Sample characteristics as well as geographic position can be found in S1 table.

### DNA extraction

HT DNA extractions were performed in 1.2 mL 2D barcoded matrix tubes pre-filled with lysing matrix E (1.4 mm ceramic spheres, 0.1 mm silica spheres, and one 4 mm glass bead) from MP biomedicals (https://www.mpbio.com/bs/116984001b-lysing-matrix-e-barcoded-plate). Lysing Matrix E has previously been shown to effectively lyse both gram-positive and negative bacteria (51). Before extraction, 100 µL (∼125 mg soil) of each soil type was transferred to a 2D barcoded Lysing Matrix E tube with a 1 mL syringe, whereafter the sample barcode was linked to the 2D extraction tube barcode with a Mirage Rack Reader (Ziath) and the software DataPaq™ (Ziath). DNA extractions with FastDNA™ SPIN kit for Soil and DNeasy® PowerSoil® Kit were performed with the available kit lysis tubes. Soil target input was 125 mg unless otherwise stated.

DNA extraction followed the manufacturer’s protocol for all kits except for DNeasy® 96 PowerSoil® Pro QIAcube® HT Kit, which followed a slightly modified protocol.

### DNeasy® 96 PowerSoil® Pro QIAcube® HT Kit

DNA extraction followed a slightly modified protocol of the DNeasy® 96 Powersoil® Pro QIAcube® HT Kit. Firstly, 500 µL CD1 was added to 125 mg of soil (unless otherwise stated), whereafter samples underwent three bead-beating cycles performed in two-minute intervals using the FastPrep96™. Between rounds of bead-beating the samples were kept on ice for two minutes. After lysis, samples were centrifuged at 3.486 x g for 10 minutes using an Eppendorf 5810 benchtop centrifuge, and 300 µL supernatant was transferred to a clean S-block containing 300 µL CD2 and 100 µL nuclease-free water (NFW) to meet the requirement of 700 µL for the rest of the protocol. Samples were again centrifuged at 3.486 x g for 10 minutes, whereafter subsequent steps followed the manufacturer’s protocol. The sample transfer step was done using the QIAcube® HT.

### V4 Amplicons: PCRBIO Ultramix

Standard amplicon libraries were prepared as one 50 µL reaction and subsequently split into two 25 µL reactions. Up to 20 ng of quality-controlled genomic DNA was used as the template. After 25 cycles of PCR (amplicon PCR) duplicate samples were pooled and cleaned using 0.8x CleanNGS sample purification beads and washed twice with 80 % EtOH and eluted in NFW. Another 8 cycles of library PCR were performed on up to 10 ng of amplicon template and cleaned as previously described. Final libraries were quantified and pooled equimolarly to produce the final sequencing libraries. Quality control was performed using the Qubit 1X HS assay [Invitrogen™, Thermo Fisher] and either Genomic DNA ScreenTape or D1000 ScreenTape [Agilent Technologies].

Miniaturized amplicon libraries were prepared with the I.DOT One as two individual 5 µL reactions by dispensing into two individual PCR plates. Each 5 µL amplicon PCR reaction consisted of 1.5 µL sample/NFW (target: 2 ng DNA), 2.5 µL PCRBIO 2x Ultra Mix, and 1 µL abV4-C tailed amplicon primer mix (2 µM, 400 nM final concentration). The subsequent 5 µL library PCR was prepared with 2 ng of cleaned PCR template in 1.5 µL/NFW (target: 2 ng DNA), 2.5 µL PCRBIO 2x Ultra Mix, and 0.5 µL adapter indexes (4 µM). After clean-up, libraries were pooled equimolarly using the I.DOT One.

A detailed protocol can be found at: https://github.com/SebastianDall/HT-downscaled-amplicon-library-protocol.

Amplicons were sequenced on the Illumina MiSeq platform. ASV abundance tables were generated by running AmpProc 5.1 (https://github.com/eyashiro/AmpProc) using the following choices: standard workflow, generate both otu and zotu tables, process only single-end reads, no primer region removal, amplicon region V4 and a version of the SILVA SSURef 99 % v138.1 database (52) processed by AutoTax (53). AmpProc is a wrapper script for running USEARCH11 (54) and downstream processing of output tables. AmpProc assigns taxonomy to ASVs by running SINTAX with the confidence cutoff set to 0.8 (55).

### Metagenomes: Illumina DNA prep

Standard metagenomic libraries were prepared according to the recommendations in the Illumina DNA prep protocol [Illumina] but eluted in NFW instead of the resuspension buffer (RSB).

Miniaturized metagenomic libraries were prepared with the I.DOT One and followed a 1:10 reagent volume reduction of the Illumina DNA prep protocol. Firstly, a 3 µL template (target: 20 ng DNA) was prepared with the I.DOT One, whereafter 2 µL BLT/TB1 master mix was added using the I.DOT One. The reaction was incubated in a thermocycler running the TAG-program from the Illumina DNA Prep protocol. The tagmentation reaction was stopped by adding 1 µL TSB using the I.DOT One, and incubation of the reaction with the PTC program in a thermocycler. After stopping the tagmentation, the libraries were washed twice with 10 µL TWB. 2 µL EPM, 2 µL NFW and 1 µL IDT® Illumina UD index was added to each well by the I.DOT One and using an epMotion® 96 [Eppendorf], respectively. Based on the original genomic DNA input, libraries were given 7 (>=4.9 ng), 8 (2.5-4.9 ng), 10 (0.9-2.5 ng), or 14 (<0.9 ng) cycles of the BLT-PCR program. PCR reactions were diluted with 17 µL NFW before 18 µL of the reactions were transferred to a new PCR-plate with 16 µL of sample purification beads (SPB) and 18 µL NFW in each well. After incubation, the beads were allowed to pellet before 50 µL of the supernatant was transferred to a new PCR-plate with 6 µL of SPB. After another incubation step, the beads were washed twice with 45 µL of 80 % ethanol and subsequently eluted in 20 µL NFW.

Final libraries were quantified and pooled equimolarly to produce the final sequencing libraries. Quality control was performed using the Qubit 1X HS assay [Invitrogen™, Thermo Fisher] and DS1000 or DS1000 HS ScreenTape [Agilent Technologies]. A detailed protocol can be found at https://github.com/SebastianDall/HT-Downscaled-Illumina-Metagenomes-Protocol.

Metagenome libraries were sequenced with the Illumina NovaSeq 6000 to a median depth of 5 Gbp. Raw Illumina reads were trimmed for barcodes, quality filtered, and deduplicated with fastp (56) and GNU-parallel (57), whereafter 16S rRNA genes were extracted from the quality-filtered reads. To extract the 16S reads, HMM models of the Rfam 14.7 seed alignments for Bacteria (RF00177) and Archaea (RF01959) were built (58) with hmmbuild (HMMER V3.3.2) (59). The HMM models were used with nhmmer and seqkit (60) to extract the 16S reads which were quality filtered for the best match using the bit-score. The 16S reads were taxonomically assigned using SINTAX with the confidence cutoff set to 0.8 using the previously mentioned database. The output was transformed into an observational table and aggregated to the genus level using R. The scripts and parameters can be found at https://github.com/SebastianDall/MFD_HT_PAPER.

### Visualization and Statistical Analysis

All data visualization and statistical analysis were carried out in R (4.2) and Rstudio (2023.03.0+386) with the following packages: tidyverse (61), ampvis2 (62), vegan (63), and bioconductor (64). Source code can be found at https://github.com/SebastianDall/MFD_HT_PAPER.

## Data Availability Statement

All sequencing data have been deposited in the ENA database under accession number PRJEB65366. Scripts for processing data and generating figures can be found on GitHub https://github.com/SebastianDall/MFD_HT_PAPER.

## Author Contributions

Conceived and designed the experiments: TBNJ SMD SMK SK MA. Performed the experiments: TBNJ SMD SK. Analyzed the data: TBNJ SMD SK SMK MA. Contributed reagents/materials/analysis tools: TBNJ SMD SMK MA. Wrote the paper: TBNJ SMD MA.

---

**S1 Fig.**
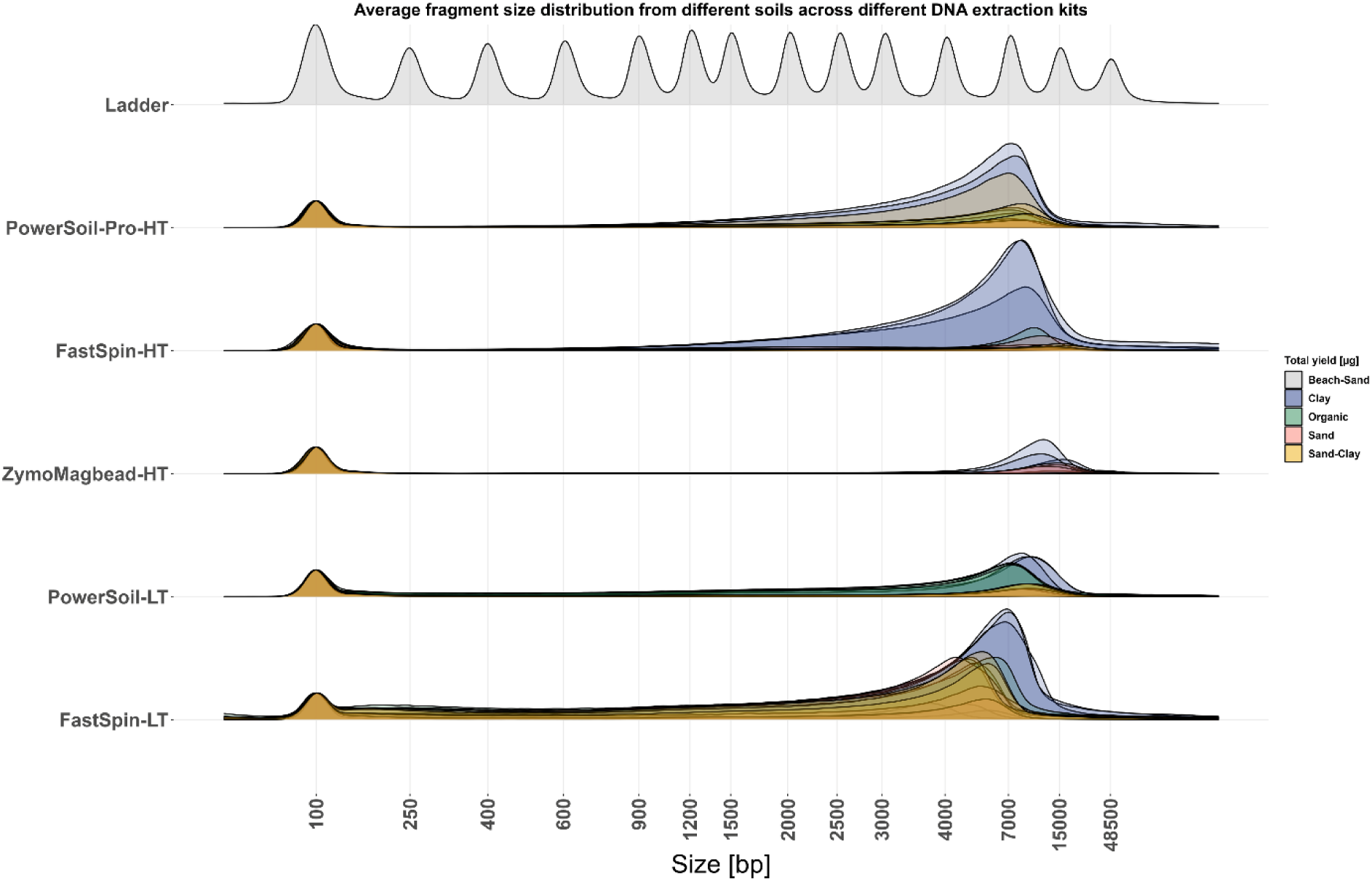
Genomic DNA fragment distribution across different DNA extraction kits. Each replicate is colored by soil type.

**S2 Fig.**
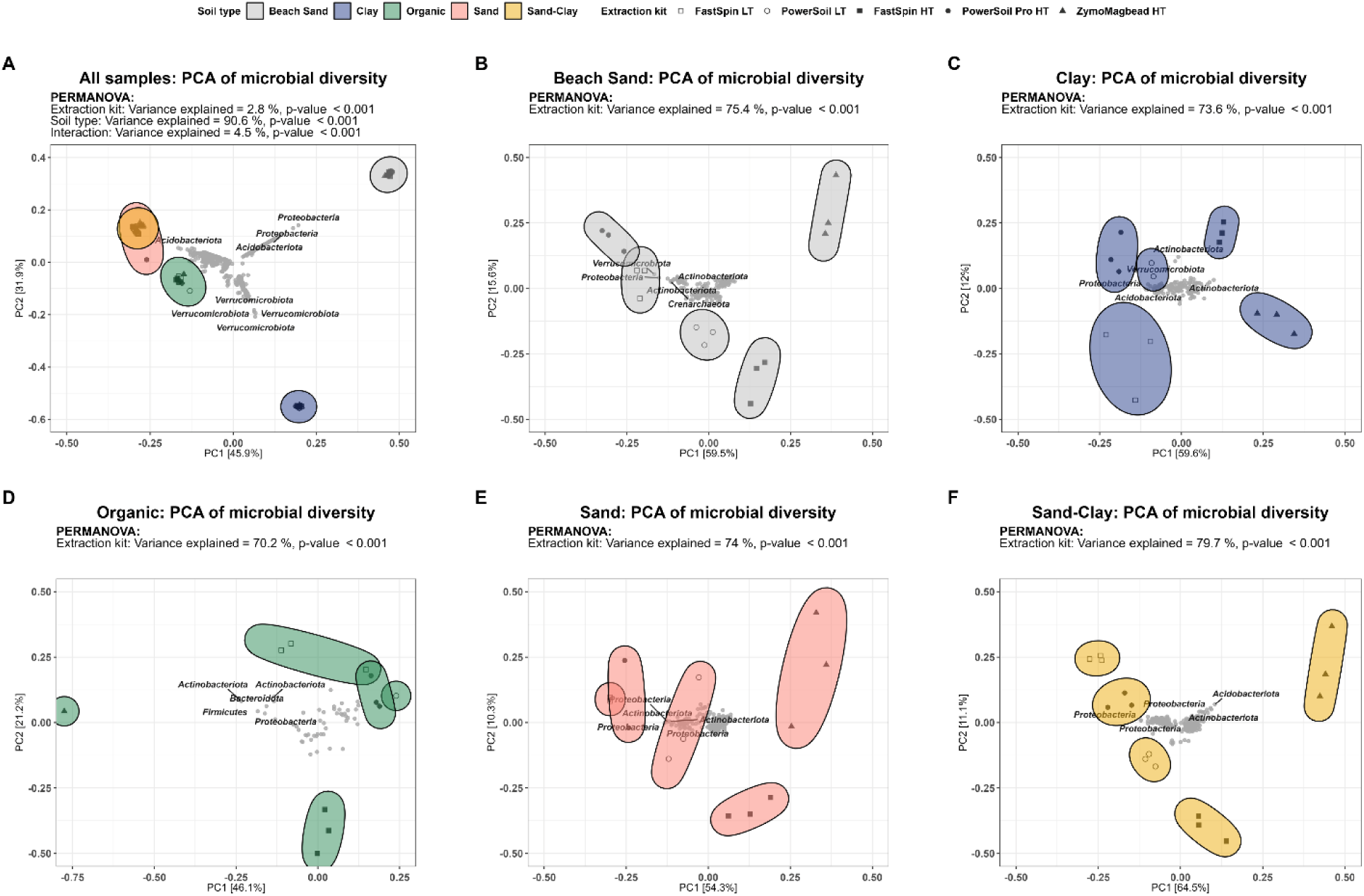
PCA of DNA extraction kits stratified by soil type. (A) Beach Sand, (B) Clay, (C) Organic, (D) Sand, (E) Sand-Clay. OTUs not xceeding 0.1% relative abundance in at least one sample were removed before Hellinger-transformation.

**S3 Fig.**
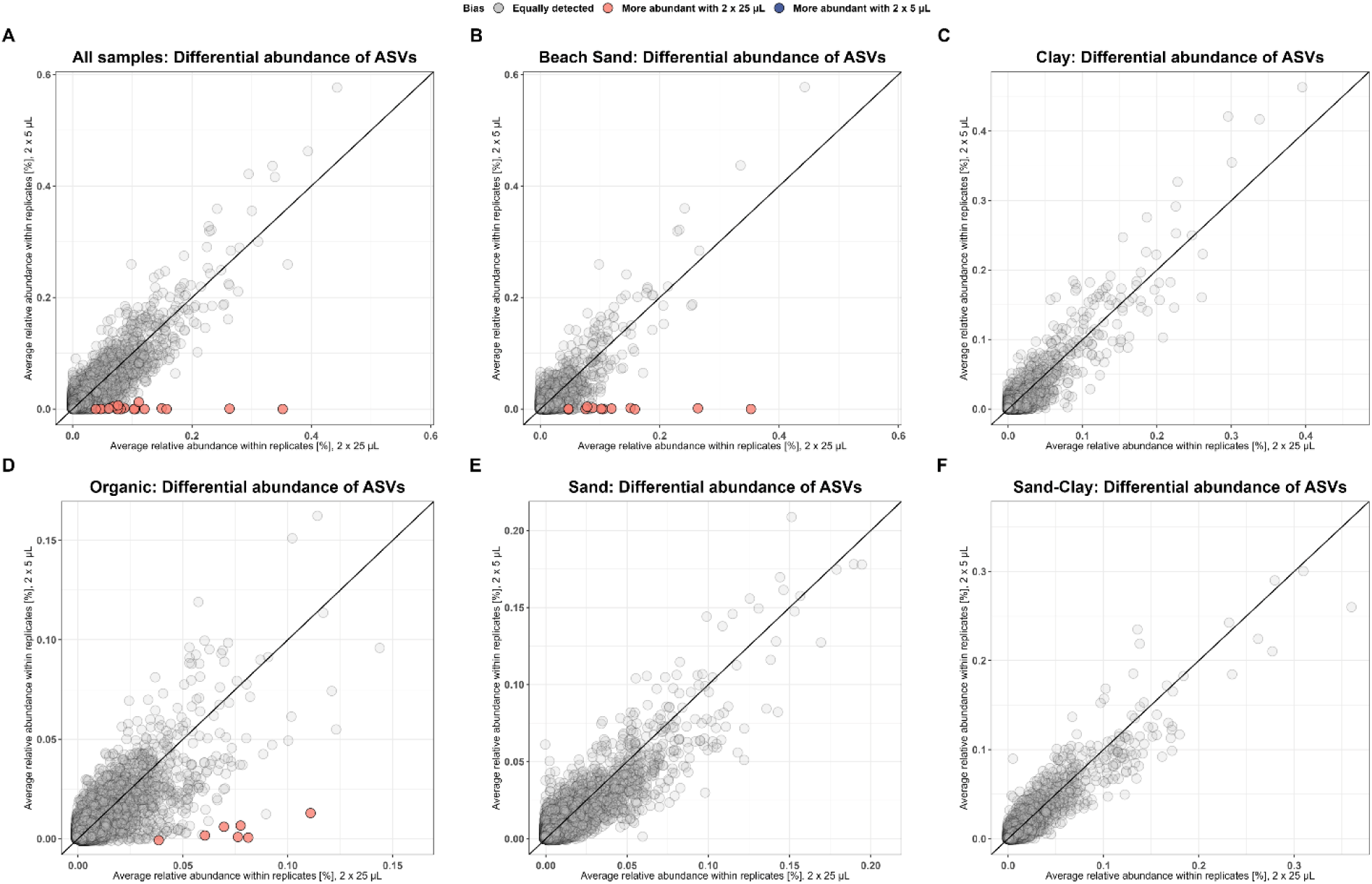
Differential abundance plot for each soil type. (A) Differential abundance plot for each soil type: (B) Beach Sand, (C) Clay, (D) rganic, (E) Sand, (F) Sand-Clay. ASVs differentially abundant in one protocol are marked with color. Bias was calculated with DESeq2 nd is defined as a significant difference in log2-fold-change (adjusted p-value<0.05).

**S4 Fig.**
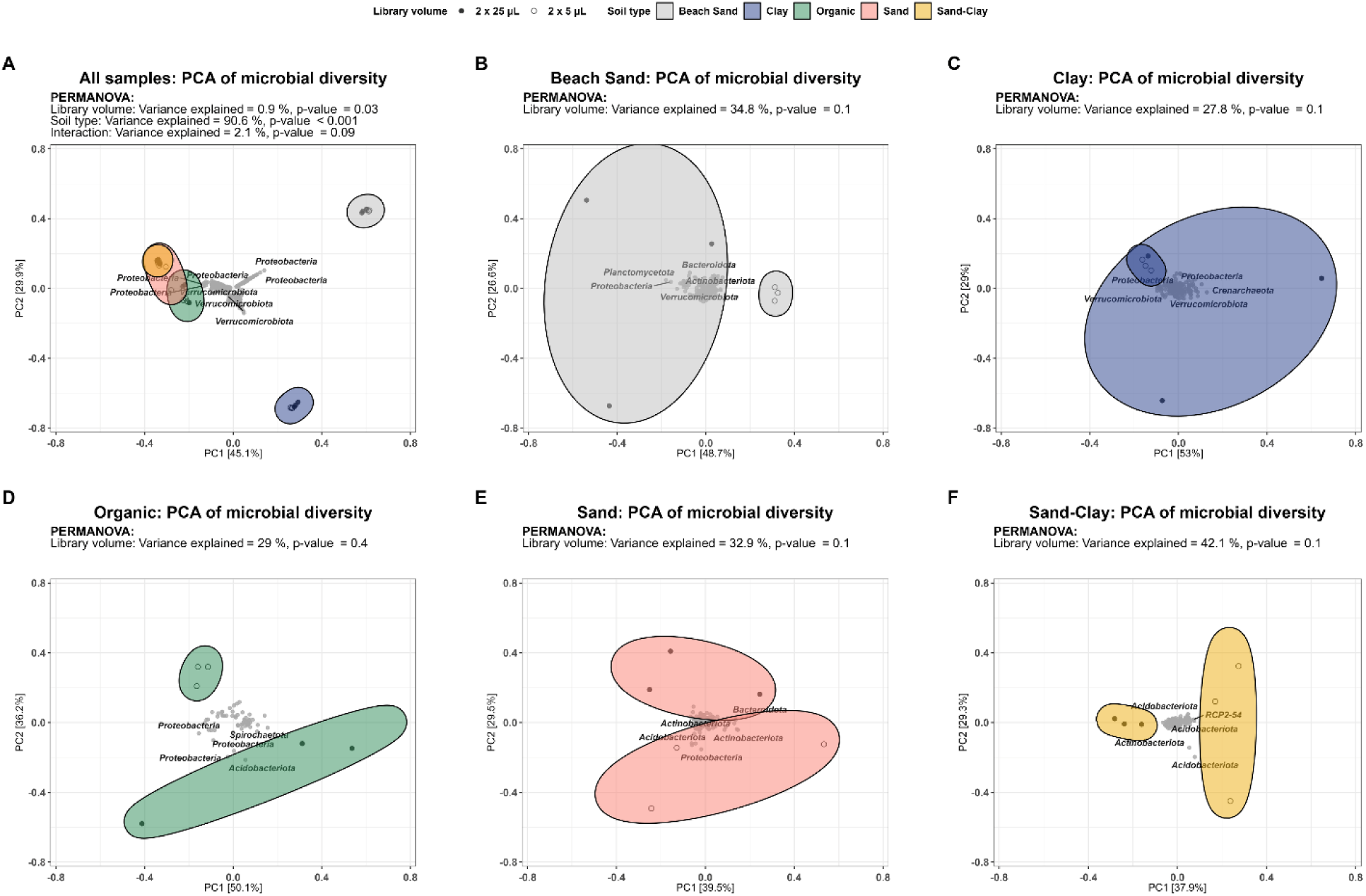
PCA of the miniaturized and standard protocol for amplicons. (A) PCA of miniaturized and standard amplicon protocol. PCA f miniaturized and standard amplicon protocol stratified by soil type: (B) Beach Sand, (C) Clay, (D) Organic, (E) Sand, (F) Sand-Clay. SVs not exceeding 0.1% relative abundance in at least one sample were removed before Hellinger-transformation.

**S5 Fig.**
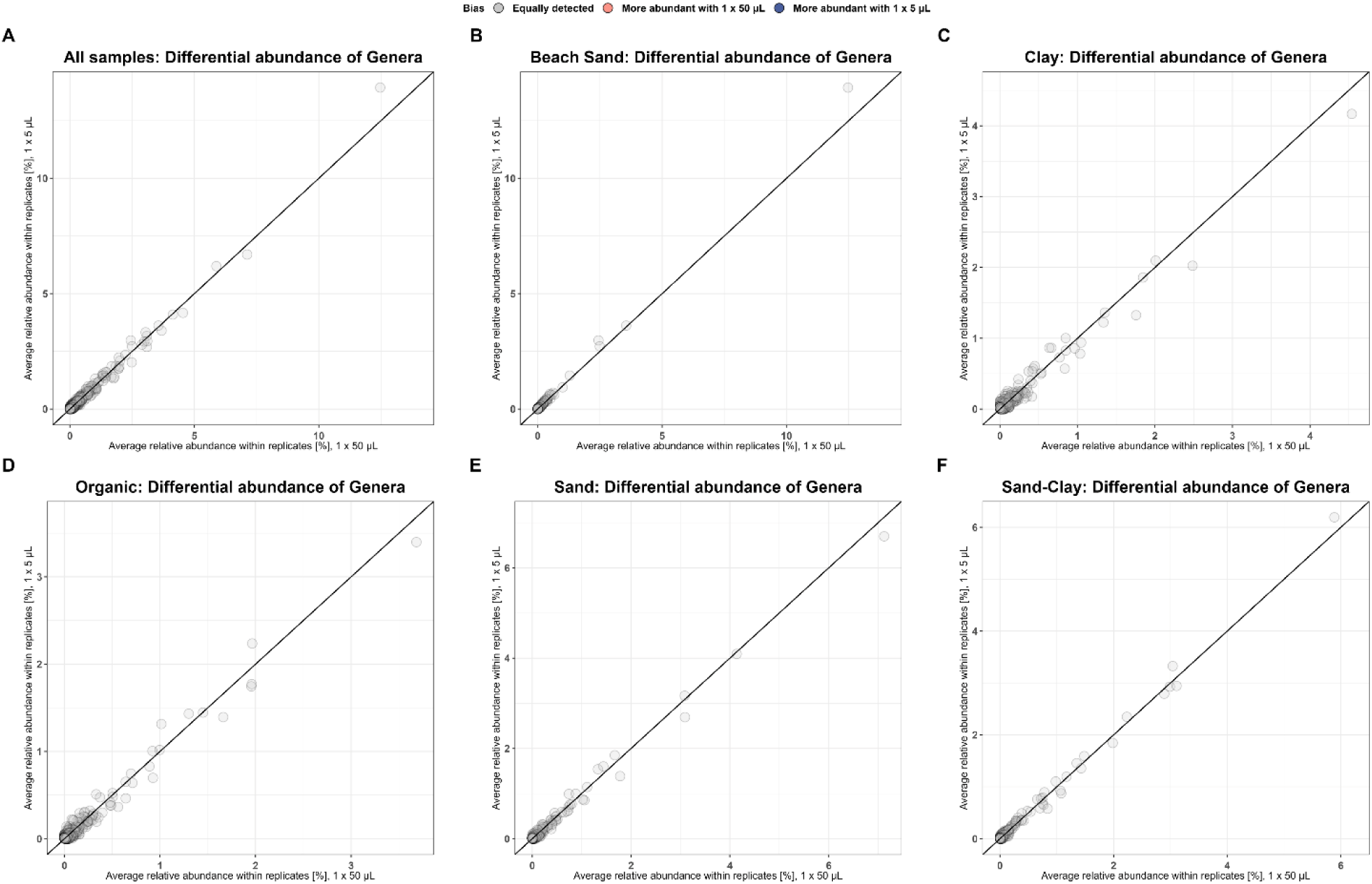
Differential abundance plots comparing the miniaturized and standard Illumina protocol. (A) Differential abundance plot for ach soil type: (B) Beach Sand, (C) Clay, (D) Organic, (E) Sand, (F) Sand-Clay. Bias was calculated with DESeq2 and is defined as a ignificant difference in log2-fold-change (adjusted p-value<0.05).

**S6 Fig.**
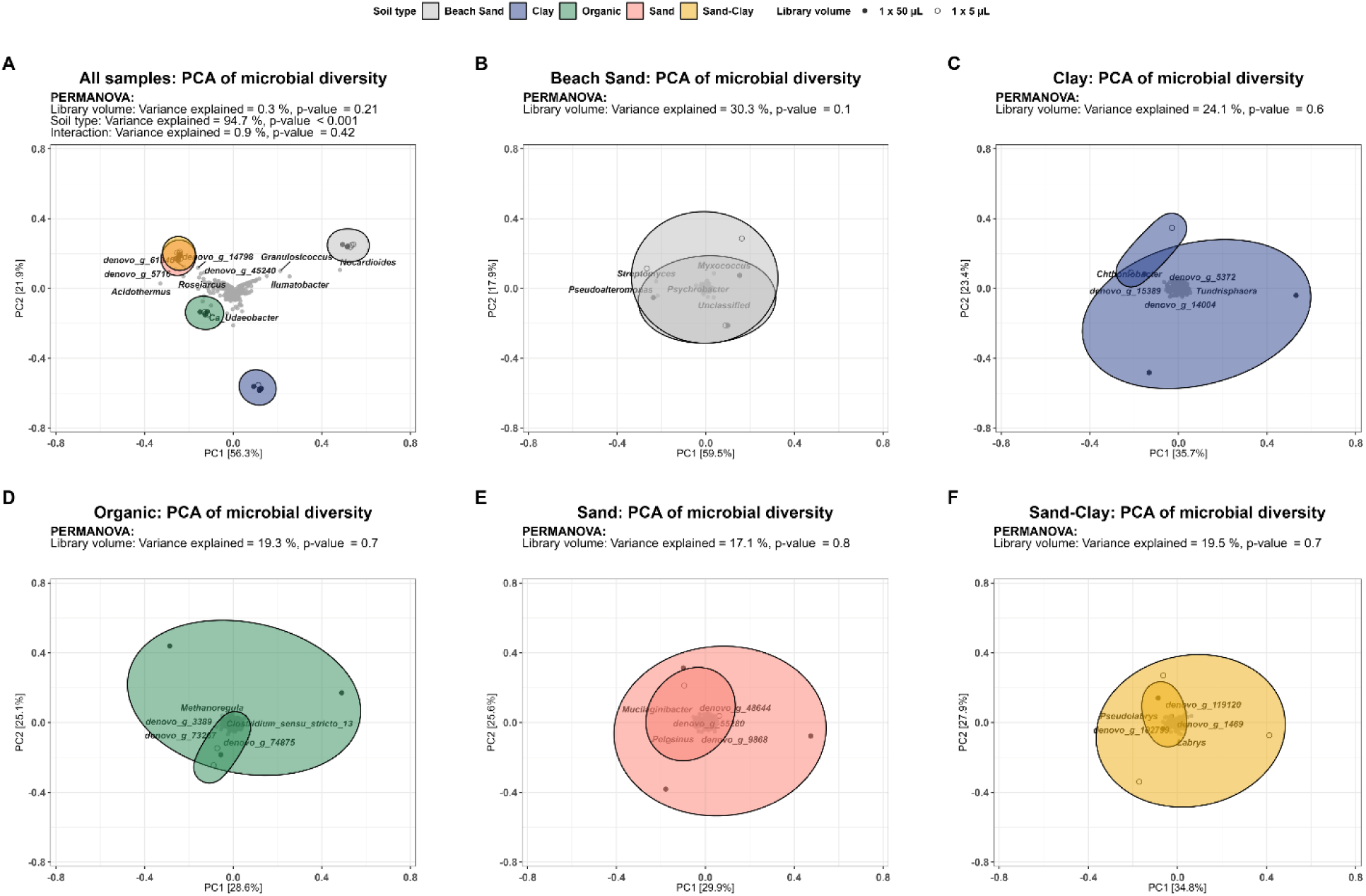
PCA of miniaturized and standard Illumina protocol. (A) PCA of miniaturized and standard Illumina protocol. PCA of iniaturized and standard illumina protocol stratified by soil type: (B) Beach Sand, (C) Clay, (D) Organic, (E) Sand, (F) Sand-Clay. Genera ot exceeding 0.1% relative abundance in at least one sample were removed before Hellinger-transformation.

**S1 Table.** Sample characteristics. Soil sample characteristics as well as location of sample site.

| Sample | Site name | Subsamples | Natura 2000 | Habitat | Longitude, Latitude | pH | Humic load |
| --- | --- | --- | --- | --- | --- | --- | --- |
| Organic | Gravlev Kær | 5 | 7230 | Alkaline fens | 56.8330, 9.8202 | 4.44 | Extreme |
| Sand | Urskoven | 5 | 9110/9120 | Beech forests | 56.8154, 9.8359 | 4.04 | High |
| Sand-Clay | Rold | 5 | 9110/9120 | Beech plantation | 56.8138, 9.8510 | 3.68 | High |
| Clay | Hestehaven | 5 | 9130 | Beech forests | 56.2853, 10.4733 | 5.80 | Intermediate |
| Beach<br>Sand | Kalø | 5 | 1220 | Perennial vegetation<br>of stony banks | 56.2732, 10.4699 | 8.90 | Low |

**S2 Table.** General statistics from DNA extraction, library preparation, and community profiles based on 16S rRNA amplicon data. All samples were rarefied to 11,468 reads (the lowest read count in any sample with more than 10,000 total reads). ASVs not exceeding 0.1 % relative abundance in at least one sample were removed prior to Hellinger-transformation and calculation of Bray-Curtis dissimilarity. Numbers represent mean and numbers in parentheses represent standard deviation. *DNA concentration could not be determined due to interference from humic substances.

| Soil type | Extraction Kit | Laboratory metrics (n=3) |  |  |  | Sequencing Metrics (n=Libraries) |  |  |  |  |  |
| --- | --- | --- | --- | --- | --- | --- | --- | --- | --- | --- | --- |
|  |  | DNA yield [µg] | 260/280 | 260/230 | Integrity [kbp] | Libraries | Library conc. [ng/µL] | Number of reads | Observed ASVs | Shannon Diversity | Bray-Curtis Dissimilarity |
| Beach Sand | FastSpin LT | NA* | 1.90 (0.09) | 0.02 (0.00) | 4.78 (1.00) | 3 | 14.29 (11.18) | 37941 (17752) | 5931 ( 15) | 8.26 (0.02) | 0.08 (0.01) |
|  | PowerSoil LT | 0.09 (0.01) | 1.48 (0.18) | 0.70 (0.19) | 10.69 (0.71) | 3 | 25.96 ( 2.62) | 44685 (11229) | 5739 (211) | 8.22 (0.11) | 0.12 (0.03) |
|  | FastSpin HT | 0.02 (0.00) | 3.17 (0.43) | 0.18 (0.01) | 13.76 (1.14) | 3 | 27.57 ( 2.08) | 49568 ( 5044) | 5918 ( 32) | 8.29 (0.02) | 0.08 (0.00) |
|  | PowerSoil Pro HT | 0.16 (0.03) | 1.79 (0.17) | 0.95 (0.04) | 7.22 (0.95) | 3 | 10.15 ( 0.97) | 39084 (10378) | 5498 ( 64) | 8.18 (0.01) | 0.11 (0.01) |
|  | ZymoMagbead HT | 0.11 (0.02) | 1.44 (0.31) | 0.03 (0.02) | 21.09 (5.60) | 3 | 19.45 ( 1.53) | 44464 (14380) | 5604 (175) | 8.22 (0.04) | 0.10 (0.02) |
| Clay | FastSpin LT | NA* | 1.75 (0.04) | 0.26 (0.01) | 6.92 (0.11) | 2 | 12.76 (11.46) | 46242 ( 5127) | 6496 ( 70) | 8.31 (0.03) | 0.07 ( NA) |
|  | PowerSoil LT | 2.99 (0.23) | 1.88 (0.01) | 1.95 (0.01) | 8.68 (0.71) | 3 | 20.30 ( 0.98) | 46931 (12289) | 6903 ( 88) | 8.46 (0.03) | 0.09 (0.01) |
|  | FastSpin HT | 2.91 (0.61) | 2.08 (0.02) | 0.67 (0.03) | 8.76 (0.40) | 3 | 24.89 ( 2.23) | 36345 (17564) | 6386 (132) | 8.27 (0.05) | 0.09 (0.01) |
|  | PowerSoil Pro HT | 4.20 (2.02) | 1.92 (0.01) | 2.15 (0.02) | 6.87 (0.32) | 3 | 18.46 ( 2.46) | 41219 (13895) | 6665 ( 67) | 8.36 (0.02) | 0.07 (0.00) |
|  | ZymoMagbead HT | 1.08 (0.27) | 1.65 (0.17) | 0.07 (0.06) | 13.00 (2.26) | 3 | 22.68 ( 1.05) | 40117 ( 7998) | 6662 (166) | 8.39 (0.04) | 0.09 (0.00) |
| Organic | FastSpin LT | NA* | 1.56 (0.02) | 0.38 (0.07) | 5.45 (0.67) | 1 | 5.94 (10.19) | 35883 ( NA) | 8008 ( NA) | 8.76 ( NA) | NA ( NA) |
|  | PowerSoil LT | 2.82 (0.22) | 1.87 (0.01) | 1.89 (0.03) | 6.78 (0.06) | 3 | 20.75 ( 1.38) | 54391 (11029) | 8281 ( 49) | 8.81 (0.01) | 0.13 (0.01) |
|  | FastSpin HT | 1.08 (0.33) | 2.88 (0.48) | 0.21 (0.08) | 17.16 (5.66) | 3 | 16.88 (11.17) | 40582 (11119) | 7600 (356) | 8.70 (0.06) | 0.18 (0.01) |
|  | PowerSoil Pro HT | 1.75 (0.10) | 1.93 (0.02) | 1.80 (0.36) | 7.04 (0.23) | 3 | 15.51 ( 2.60) | 37852 ( 5176) | 8117 (155) | 8.78 (0.04) | 0.16 (0.01) |
|  | ZymoMagbead HT | 0.07 (0.07) | 1.55 (0.24) | 0.06 (0.01) | 11.18 (2.56) | 1 | 7.20 (12.40) | 51355 ( NA) | 7455 ( NA) | 8.64 ( NA) | NA ( NA) |
| Sand | FastSpin LT | NA* | 1.62 (0.04) | 0.14 (0.04) | 5.24 (0.69) | 3 | 17.94 ( 1.27) | 35966 (12534) | 6885 ( 15) | 8.52 (0.01) | 0.11 (0.01) |
|  | PowerSoil LT | 0.77 (0.02) | 1.78 (0.06) | 1.43 (0.18) | 8.31 (0.60) | 3 | 21.90 ( 2.18) | 34231 (11585) | 6802 (112) | 8.49 (0.03) | 0.10 (0.01) |
|  | FastSpin HT | 0.71 (0.21) | 2.65 (0.26) | 0.26 (0.04) | 11.09 (0.82) | 2 | 23.48 ( 0.72) | 44452 ( 1237) | 6447 ( 16) | 8.40 (0.02) | 0.09 ( NA) |
|  | PowerSoil Pro HT | 1.38 (0.26) | 1.91 (0.01) | 1.59 (0.38) | 7.59 (0.63) | 2 | 19.84 ( 6.02) | 48816 ( 2551) | 6798 (107) | 8.50 (0.02) | 0.13 ( NA) |
|  | ZymoMagbead HT | 0.41 (0.15) | 1.68 (0.10) | 0.03 (0.01) | 13.65 (1.16) | 3 | 21.04 ( 3.39) | 42188 ( 8694) | 6396 ( 88) | 8.37 (0.03) | 0.10 (0.01) |
| Sand-Clay | FastSpin LT | NA* | 1.53 (0.03) | 0.26 (0.04) | 5.45 (0.41) | 3 | 18.60 ( 2.10) | 45471 ( 5312) | 6710 ( 74) | 8.46 (0.01) | 0.08 (0.00) |
|  | PowerSoil LT | 1.09 (0.05) | 1.78 (0.03) | 1.48 (0.11) | 8.68 (0.20) | 3 | 19.42 ( 2.04) | 43166 (10862) | 6584 ( 34) | 8.43 (0.01) | 0.08 (0.00) |
|  | FastSpin HT | 0.80 (0.09) | 2.51 (0.10) | 0.23 (0.02) | 15.39 (2.51) | 3 | 26.02 ( 2.07) | 46872 ( 8983) | 6185 ( 55) | 8.30 (0.01) | 0.08 (0.01) |
|  | PowerSoil Pro HT | 1.25 (0.16) | 1.90 (0.04) | 1.67 (0.13) | 7.76 (0.93) | 3 | 15.66 ( 0.74) | 32527 (17196) | 6506 (173) | 8.39 (0.04) | 0.10 (0.02) |
|  | ZymoMagbead HT | 0.03 (0.01) | 1.38 (0.12) | 0.03 (0.01) | 21.28 (4.34) | 3 | 11.14 ( 7.10) | 28814 ( 2419) | 6096 (139) | 8.31 (0.05) | 0.13 (0.00) |

**S3 Table.** General library characteristics of the miniaturized and standard amplicon protocol. Microbial community characteristics of the standard and miniaturized amplicon library protocol for five different soil. All samples were rarefied to 3,511 reads (the lowest read count in any sample with more than 3,000 total reads). ASVs not exceeding 0.1 % relative abundance in at least one sample were removed prior to Hellinger-transformation and calculation of Bray-Curtis dissimilarity. Numbers represent mean (n=”Libraries”) and numbers in parentheses represent standard deviation. The number of comparisons for “Bray-Curtis dissimilarity between protocols” was 9.

| Soil type | Protocol | Libraries | Library conc.<br>[ng/μL] | Read Count | Observed<br>ASVs | Shannon<br>Diversity | Bray-Curtis<br>Dissimilarity | BC Dissimilarity<br>Between<br>Protocols |
| --- | --- | --- | --- | --- | --- | --- | --- | --- |
| Beach Sand | 2 x 25 μL | 3 | 3.92 (1.20) | 4314 ( 834) | 2394 (15) | 7.60 (0.02) | 0.29 (0.00) | 0.34 (0.02) |
| | 2 x 5 μL | 3 | 10.15 (0.97) | 39084 (10378) | 2368 (27) | 7.57 (0.02) | 0.30 (0.00) | |
| Clay | 2 x 25 μL | 3 | 8.84 (7.48) | 7414 ( 5359) | 2538 (65) | 7.65 (0.04) | 0.30 (0.01) | 0.30 (0.02) |
| | 2 x 5 μL | 3 | 18.46 (2.46) | 41212 (13895) | 2499 (15) | 7.61 (0.01) | 0.28 (0.01) | |
| Organic | 2 x 25 μL | 3 | 14.69 (2.23) | 17742 ( 2182) | 2798 (96) | 7.83 (0.05) | 0.48 (0.07) | 0.48 (0.03) |
| | 2 x 5 μL | 3 | 15.51 (2.60) | 37851 ( 5175) | 2919 (35) | 7.90 (0.02) | 0.46 (0.03) | |
| Sand | 2 x 25 μL | 3 | 12.65 (1.11) | 18601 ( 2987) | 2566 (70) | 7.71 (0.04) | 0.34 (0.01) | 0.36 (0.01) |
| | 2 x 5 μL | 3 | 19.84 (6.02) | 36365 (21638) | 2672 (38) | 7.76 (0.02) | 0.36 (0.03) | |
| Sand-Clay | 2 x 25 μL | 3 | 10.52 (1.56) | 18993 ( 2423) | 2524 (33) | 7.66 (0.02) | 0.30 (0.01) | 0.33 (0.01) |
| | 2 x 5 μL | 3 | 15.66 (0.74) | 32526 (17195) | 2528 (76) | 7.67 (0.05) | 0.33 (0.02) | |

**S4 Table.** General library characteristics of the miniaturized and standard metagenome protocol. Microbial community characteristics of the standard and miniaturized metagenome library protocol for five different soil. All samples were rarefied to 3,665 reads (lowest read count in any sample with more than 3,000 total reads). Genera not exceeding 0.1 % relative abundance in at least one sample were removed prior to Hellinger transformation and calculation of Bray-Curtis dissimilarity. Numbers represent mean (n=”Libraries”) and numbers in parentheses represent standard deviation. Number of comparisons for “Bray-Curtis dissimilarity between protocols” were 9 except for Clay (n=6).

| Soil type | Protocol | Libraries | Library conc.<br>[ng/μL] | Total Reads<br>[Gbp] | Trimmed Reads<br>[Gbp] | 16S Reads<br>Count | Observed<br>OTUs | Shannon<br>Diversity | Bray-Curtis<br>Dissimilarity | BC Dissimilarity<br>Between Protocols |
| --- | --- | --- | --- | --- | --- | --- | --- | --- | --- | --- |
| Beach<br>Sand | 1 x 50 μL | 3 | 21.43 (2.38) | 24.25 (5.09) | 12.52 (2.87) | 31527 (6788) | 312 (30) | 2.62 (0.05) | 0.22 (0.02) | 0.20 (0.02) |
| | 1 x 5 μL | 3 | 1.47 (0.53) | 16.50 (3.27) | 9.65 (1.71) | 31414 (6864) | 311 ( 1) | 2.78 (0.05) | 0.20 (0.02) | |
| Clay | 1 x 50 μL | 3 | 7.23 (2.67) | 4.28 (0.92) | 2.93 (0.64) | 4853 ( 974) | 309 (26) | 3.23 (0.02) | 0.26 (0.01) | 0.25 (0.02) |
| | 1 x 5 μL | 2 | 0.57 (0.07) | 4.18 (0.05) | 2.91 (0.05) | 5435 ( 24) | 320 ( 3) | 3.17 (0.04) | 0.24 (NA) | |
| Organic | 1 x 50 μL | 3 | 9.32 (3.45) | 3.78 (0.96) | 2.56 (0.66) | 5283 (1187) | 286 (17) | 2.99 (0.03) | 0.26 (0.01) | 0.25 (0.01) |
| | 1 x 5 μL | 3 | 0.76 (0.24) | 5.03 (1.09) | 3.41 (0.72) | 8160 (1477) | 310 ( 5) | 2.96 (0.05) | 0.23 (0.02) | |
| Sand | 1 x 50 μL | 3 | 10.13 (4.49) | 4.10 (1.07) | 2.81 (0.70) | 5042 (1322) | 189 (23) | 2.87 (0.07) | 0.22 (0.01) | 0.21 (0.02) |
| | 1 x 5 μL | 3 | 0.68 (0.13) | 5.73 (1.77) | 3.85 (1.26) | 7576 (2566) | 202 ( 5) | 2.77 (0.06) | 0.21 (0.01) | |
| Sand-Clay | 1 x 50 μL | 3 | 19.80 (1.45) | 11.03 (2.40) | 7.87 (1.74) | 13077 (2487) | 192 ( 4) | 2.77 (0.05) | 0.19 (0.00) | 0.19 (0.02) |
| | 1 x 5 μL | 3 | 0.28 (0.02) | 4.84 (1.08) | 3.36 (0.77) | 6016 (1002) | 193 ( 7) | 2.88 (0.05) | 0.20 (0.01) | |

## Bibliography

1. Bollmann-Giolai A, Giolai M, Heavens D, Macaulay I, Malone J, Clark MD. A low-cost pipeline for soil microbiome profiling. Microbiologyopen. 2020 Dec;9(12):e1133.

2. Curtis T. Microbial ecologists: it’s time to “go large.” Nat Rev Microbiol. 2006 Jul;4(7):488.

3. Thompson LR, Sanders JG, McDonald D, Amir A, Ladau J, Locey KJ, et al. A communal catalogue reveals Earth’s multiscale microbial diversity. Nature. 2017 Nov 23;551(7681):457–63.

4. Nesme J, Achouak W, Agathos SN, Bailey M, Baldrian P, Brunel D, et al. Back to the Future of Soil Metagenomics. Front Microbiol. 2016 Feb 10;7:73.

5. Marotz C, Amir A, Humphrey G, Gaffney J, Gogul G, Knight R. DNA extraction for streamlined metagenomics of diverse environmental samples. Biotechniques. 2017 Jun 1;62(6):290–3.

6. Mayday MY, Khan LM, Chow ED, Zinter MS, DeRisi JL. Miniaturization and optimization of 384-well compatible RNA sequencing library preparation. PLoS One. 2019 Jan 10;14(1):e0206194.

7. Minich JJ, Humphrey G, Benitez RAS, Sanders J, Swafford A, Allen EE, et al. High-Throughput Miniaturized 16S rRNA Amplicon Library Preparation Reduces Costs while Preserving Microbiome Integrity. mSystems [Internet]. 2018 Nov 6;3(6). Available from: 10.1128/mSystems.00166-18

8. Mora-Castilla S, To C, Vaezeslami S, Morey R, Srinivasan S, Chousal JN, et al. Miniaturization Technologies for Efficient Single-Cell Library Preparation for Next-Generation Sequencing. J Lab Autom. 2016 Aug;21(4):557–67.

9. Fornasier F, Ascher J, Ceccherini MT, Tomat E, Pietramellara G. A simplified rapid, low-cost and versatile DNA-based assessment of soil microbial biomass. Ecol Indic. 2014 Oct 1;45:75–82.

10. Chiodi C, Moro M, Squartini A, Concheri G, Occhi F, Fornasier F, et al. High-Throughput Isolation of Nucleic Acids from Soil. Soil Systems. 2019 Dec 28;4(1):3.

11. Li T, Zhang S, Hu J, Hou H, Li K, Fan Q, et al. Soil sample sizes for DNA extraction substantially affect the examination of microbial diversity and co-occurrence patterns but not abundance. Soil Biol Biochem. 2023 Feb 1;177:108902.

12. Habtom H, Demanèche S, Dawson L, Azulay C, Matan O, Robe P, et al. Soil characterisation by bacterial community analysis for forensic applications: A quantitative comparison of environmental technologies. Forensic Sci Int Genet. 2017 Jan;26:21–9.

13. Feinstein LM, Sul WJ, Blackwood CB. Assessment of bias associated with incomplete extraction of microbial DNA from soil. Appl Environ Microbiol. 2009 Aug;75(16):5428– 33.

14. Human Microbiome Project Consortium. Structure, function and diversity of the healthy human microbiome. Nature. 2012 Jun 13;486(7402):207–14.

15. Cowan DA, Lebre PH, Amon C, Becker RW, Boga HI, Boulangé A, et al. Biogeographical survey of soil microbiomes across sub-Saharan Africa: structure, drivers, and predicted climate-driven changes. Microbiome. 2022 Aug 23;10(1):131.

16. Iturbe-Espinoza P, Brandt BW, Braster M, Bonte M, Brown DM, van Spanning RJM. Effects of DNA preservation solution and DNA extraction methods on microbial community profiling of soil. Folia Microbiol . 2021 Aug;66(4):597–606.

17. Hermans SM, Buckley HL, Lear G. Optimal extraction methods for the simultaneous analysis of DNA from diverse organisms and sample types. Mol Ecol Resour. 2018 May;18(3):557–69.

18. Liu D, Pérez-Moreno J, He X, Garibay-Orijel R, Yu F. Truffle Microbiome Is Driven by Fruit Body Compartmentalization Rather than Soils Conditioned by Different Host Trees. mSphere. 2021 Aug 25;6(4):e0003921.

19. Saidi-Mehrabad A, Neuberger P, Cavaco M, Froese D, Lanoil B. Optimization of subsampling, decontamination, and DNA extraction of difficult peat and silt permafrost samples. Sci Rep. 2020 Aug 31;10(1):14295.

20. Knudsen BE, Bergmark L, Munk P, Lukjancenko O, Priemé A, Aarestrup FM, et al. Impact of Sample Type and DNA Isolation Procedure on Genomic Inference of Microbiome Composition. mSystems [Internet]. 2016 Oct 18;1(5). Available from: 10.1128/mSystems.00095-16

21. Young JM, Rawlence NJ, Weyrich LS, Cooper A. Limitations and recommendations for successful DNA extraction from forensic soil samples: a review. Sci Justice. 2014 May;54(3):238–44.

22. Shaffer JP, Carpenter CS, Martino C, Salido RA, Minich JJ, Bryant M, et al. A comparison of six DNA extraction protocols for 16S, ITS and shotgun metagenomic sequencing of microbial communities. Biotechniques. 2022 Jun;73(1):34–46.

23. Sui H-Y, Weil AA, Nuwagira E, Qadri F, Ryan ET, Mezzari MP, et al. Impact of DNA Extraction Method on Variation in Human and Built Environment Microbial Community and Functional Profiles Assessed by Shotgun Metagenomics Sequencing. Front Microbiol. 2020 May 25;11:953.

24. Yang J, Sooksa-Nguan T, Kannan B, Cano-Alfanar S, Liu H, Kent A, et al. Microbiome differences in sugarcane and metabolically engineered oilcane accessions and their implications for bioenergy production. Biotechnol Biofuels Bioprod. 2023 Mar 30;16(1):56.

25. Farhadfar N, Gharaibeh RZ, Dahl WJ, Mead L, Alabasi KM, Newsome R, et al. Gut Microbiota Dysbiosis Associated with Persistent Fatigue in Hematopoietic Cell Transplantation Survivors. Transplant Cell Ther. 2021 Jun;27(6):498.e1–498.e8.

26. Davis Birch WA, Moura IB, Ewin DJ, Wilcox MH, Buckley AM, Culmer PR, et al. MiGut: A scalable in vitro platform for simulating the human gut microbiome-Development, validation and simulation of antibiotic-induced dysbiosis. Microb Biotechnol [Internet]. 2023 Apr 10; Available from: 10.1111/1751-7915.14259

27. Ascher J, Ceccherini MT, Pantani OL, Agnelli A, Borgogni F, Guerri G, et al. Sequential extraction and genetic fingerprinting of a forest soil metagenome. Appl Soil Ecol. 2009 Jun 1;42(2):176–81.

28. Martin-Laurent F, Philippot L, Hallet S, Chaussod R, Germon JC, Soulas G, et al. DNA extraction from soils: old bias for new microbial diversity analysis methods. Appl Environ Microbiol. 2001 May;67(5):2354–9.

29. Sansupa C, Purahong W, Wubet T, Tiansawat P, Pathom-Aree W, Teaumroong N, et al. Soil bacterial communities and their associated functions for forest restoration on a limestone mine in northern Thailand. PLoS One. 2021 Apr 8;16(4):e0248806.

30. Babalola OO, Adedayo AA, Fadiji AE. Metagenomic Survey of Tomato Rhizosphere Microbiome Using the Shotgun Approach. Microbiol Resour Announc. 2022 Feb 17;11(2):e0113121.

31. Soliman T, Yang S-Y, Yamazaki T, Jenke-Kodama H. Profiling soil microbial communities with next-generation sequencing: the influence of DNA kit selection and technician technical expertise. PeerJ. 2017 Dec 19;5:e4178.

32. Wydro U. Soil Microbiome Study Based on DNA Extraction: A Review. Water. 2022 Dec 8;14(24):3999.

33. Lori M, Armengot L, Schneider M, Schneidewind U, Bodenhausen N, Mäder P, et al. Organic management enhances soil quality and drives microbial community diversity in cocoa production systems. Sci Total Environ. 2022 Aug 15;834:155223.

34. Wang X, Reilly K, Heathcott R, Biswas A, Johnson LJ, Teasdale S, et al. Soil Nitrogen Treatment Alters Microbiome Networks Across Farm Niches. Front Microbiol. 2021;12:786156.

35. King WL, Richards SC, Kaminsky LM, Bradley BA, Kaye JP, Bell TH. Leveraging microbiome rediversification for the ecological rescue of soil function. Environ Microbiome. 2023 Jan 23;18(1):7.

36. Kaminsky LM, Esker PD, Bell TH. Abiotic conditions outweigh microbial origin during bacterial assembly in soils. Environ Microbiol. 2021 Jan;23(1):358–71.

37. Barbour KM, Weihe C, Allison SD, Martiny JBH. Bacterial community response to environmental change varies with depth in the surface soil. Soil Biol Biochem. 2022 Sep 1;172:108761.

38. McDonald MD, Lewis KL, DeLaune PB, Hux BA, Boutton TW, Gentry TJ. Nitrogen fertilizer driven nitrous and nitric oxide production is decoupled from microbial genetic potential in low carbon, semi-arid soil. Frontiers in Soil Science [Internet]. 2023;2. Available from: https://www.frontiersin.org/articles/10.3389/fsoil.2022.1050779

39. Kui L, Xiang G, Wang Y, Wang Z, Li G, Li D, et al. Large-Scale Characterization of the Soil Microbiome in Ancient Tea Plantations Using High-Throughput 16S rRNA and Internal Transcribed Spacer Amplicon Sequencing. Front Microbiol. 2021 Oct 15;12:745225.

40. Ishida JK, Bini AP, Creste S, Van Sluys M-A. Towards defining the core Saccharum microbiome: input from five genotypes. BMC Microbiol. 2022 Aug 8;22(1):193.

41. DiLegge MJ, Manter DK, Vivanco JM. Soil microbiome disruption reveals specific and general plant-bacterial relationships in three agroecosystem soils. PLoS One. 2022 Nov 16;17(11):e0277529.

42. Pan X, Angelidaki I, Alvarado-Morales M, Liu H, Liu Y, Huang X, et al. Methane production from formate, acetate and H2/CO2; focusing on kinetics and microbial characterization. Bioresour Technol. 2016 Oct;218:796–806.

43. Sato MP, Ogura Y, Nakamura K, Nishida R, Gotoh Y, Hayashi M, et al. Comparison of the sequencing bias of currently available library preparation kits for Illumina sequencing of bacterial genomes and metagenomes. DNA Res. 2019 Oct 1;26(5):391–8.

44. Marine R, Polson SW, Ravel J, Hatfull G, Russell D, Sullivan M, et al. Evaluation of a transposase protocol for rapid generation of shotgun high-throughput sequencing libraries from nanogram quantities of DNA. Appl Environ Microbiol. 2011 Nov;77(22):8071–9.

45. Thomas T, Gilbert J, Meyer F. Metagenomics - a guide from sampling to data analysis. Microb Inform Exp. 2012 Feb 9;2(1):3.

46. Gaio D, To J, Liu M, Monahan L, Anantanawat K, Darling AE. Hackflex: low cost Illumina sequencing library construction for high sample counts [Internet]. bioRxiv. 2019 [cited 2023 Mar 30]. p. 779215. Available from: https://www.biorxiv.org/content/10.1101/779215v1

47. Baym M, Kryazhimskiy S, Lieberman TD, Chung H, Desai MM, Kishony R. Inexpensive multiplexed library preparation for megabase-sized genomes. PLoS One. 2015 May 22;10(5):e0128036.

48. Koren S, Phillippy AM. One chromosome, one contig: complete microbial genomes from long-read sequencing and assembly. Curr Opin Microbiol. 2015 Feb;23:110–20.

49. Sanders JG, Yan W, Mjungu D, Lonsdorf EV, Hart JA, Sanz CM, et al. A low-cost genomics workflow enables isolate screening and strain-level analyses within microbiomes. Genome Biol. 2022 Oct 12;23(1):212.

50. Pansu M, Gautheyrou J. Handbook of Soil Analysis. Springer Berlin Heidelberg; 2006. 19 p.

51. Albertsen M, Karst SM, Ziegler AS, Kirkegaard RH, Nielsen PH. Back to Basics--The Influence of DNA Extraction and Primer Choice on Phylogenetic Analysis of Activated Sludge Communities. PLoS One. 2015 Jul 16;10(7):e0132783.

52. Quast C, Pruesse E, Yilmaz P, Gerken J, Schweer T, Yarza P, et al. The SILVA ribosomal RNA gene database project: improved data processing and web-based tools. Nucleic Acids Res. 2013 Jan;41(Database issue):D590–6.

53. Dueholm MS, Andersen KS, McIlroy SJ, Kristensen JM, Yashiro E, Karst SM, et al. Generation of Comprehensive Ecosystem-Specific Reference Databases with Species- Level Resolution by High-Throughput Full-Length 16S rRNA Gene Sequencing and Automated Taxonomy Assignment (AutoTax). MBio [Internet]. 2020 Sep 22;11(5). Available from: 10.1128/mBio.01557-20

54. Edgar RC. Search and clustering orders of magnitude faster than BLAST. Bioinformatics. 2010 Oct 1;26(19):2460–1.

55. Edgar RC. SINTAX: a simple non-Bayesian taxonomy classifier for 16S and ITS sequences [Internet]. bioRxiv. 2016 [cited 2023 May 22]. p. 074161. Available from: https://www.biorxiv.org/content/10.1101/074161v1

56. Chen S, Zhou Y, Chen Y, Gu J. fastp: an ultra-fast all-in-one FASTQ preprocessor. Bioinformatics. 2018 Sep 1;34(17):i884–90.

57. Tange, O. (2022, July 22). GNU Parallel 20220722 (’Roe vs Wade’). Zenodo. 10.5281/zenodo.6891516

58. Kalvari I, Nawrocki EP, Ontiveros-Palacios N, Argasinska J, Lamkiewicz K, Marz M, et al. Rfam 14: expanded coverage of metagenomic, viral and microRNA families. Nucleic Acids Res. 2021 Jan 8;49(D1):D192–200.

59. Wheeler TJ, Eddy SR. nhmmer: DNA homology search with profile HMMs. Bioinformatics. 2013 Oct 1;29(19):2487–9.

60. Shen W, Le S, Li Y, Hu F. SeqKit: A Cross-Platform and Ultrafast Toolkit for FASTA/Q File Manipulation. PLoS One. 2016 Oct 5;11(10):e0163962.

61. Wickham H, Averick M, Bryan J, Chang W, McGowan L, François R, et al. Welcome to the tidyverse. J Open Source Softw. 2019 Nov 21;4(43):1686.

62. Andersen KS, Kirkegaard RH, Karst SM, Albertsen M. ampvis2: an R package to analyse and visualise 16S rRNA amplicon data [Internet]. bioRxiv. 2018 [cited 2023 May 22]. p. 299537. Available from: https://www.biorxiv.org/content/10.1101/299537v1

63. Dixon P. VEGAN, a package of R functions for community ecology. J Veg Sci. 2003 Dec;14(6):927–30.

64. Huber W, Carey VJ, Gentleman R, Anders S, Carlson M, Carvalho BS, et al. Orchestrating high-throughput genomic analysis with Bioconductor. Nat Methods. 2015 Feb;12(2):115–21.

