## Supplementary material for "High-throughput DNA extraction and cost-effective miniaturized metagenome and amplicon library preparation of soil samples for DNA sequencing": S1 file

### Supplementary File 1:

#### Humic substance interference of Qubit measurements

FastDNA™ SPIN kit for soil removed less of the water-soluble part of the humic substances, which was directly observable for all soil types except Beach Sand by the brown tint of the DNA extracts. Based on the original measured concentration, a serial dilution was prepared and quantified (Fig 1). All points were fitted with a second-degree polynomial function, but a first-degree polynomial fitted the points equally well for the Qubit Standard ($R^2$=1.00, p<0.001) and the Activated sludge ($R^2$=1.00, p<0.001). As evident by the dilution series, the remaining humic substances have a direct impact on the Qubit 1X HS DNA assay leading to inaccurate DNA concentration measurements.

A

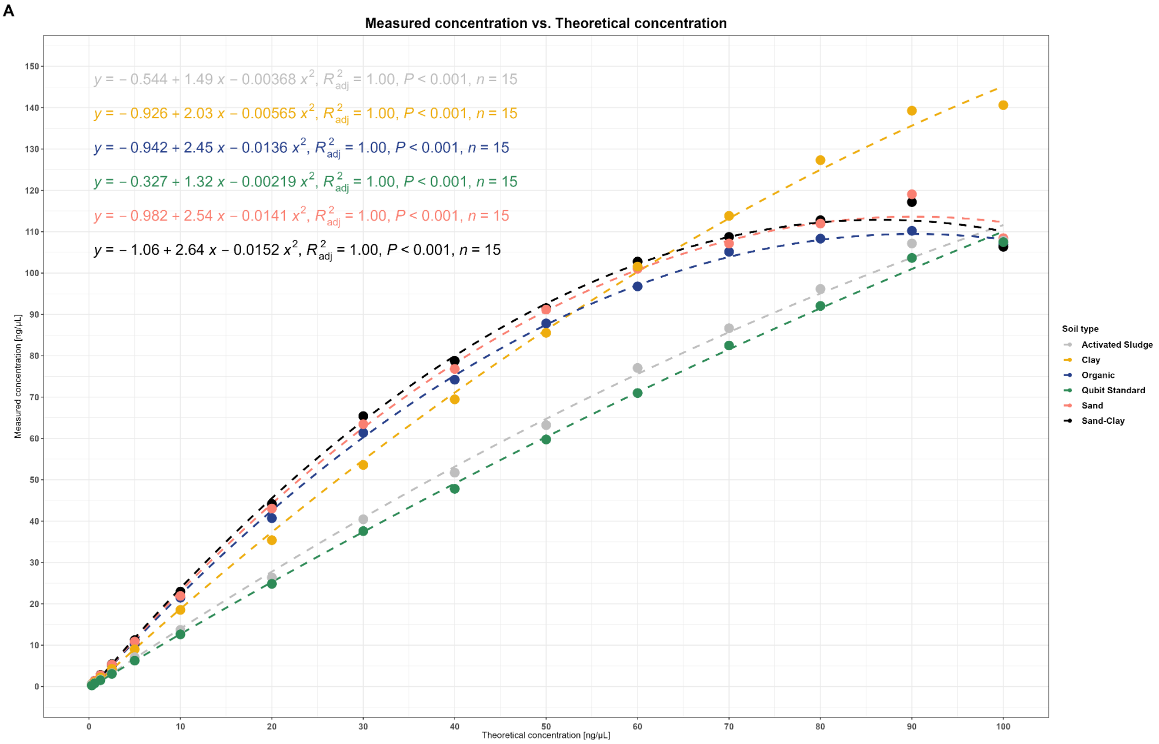


**Fig 1. Qubit DNA concentrations for soil samples processed with FastDNA SPIN kit.** Measured concentration vs. theoretical concentration in ng/µL for samples processed with FastDNA™ SPIN kit for Soil. Activated Sludge was used instead of Beach Sand as a low humic substance sample.
