## Supplementary material for "High-throughput DNA extraction and cost-effective miniaturized metagenome and amplicon library preparation of soil samples for DNA sequencing": S2 file

### Supplementary File 2:

#### Optimization of PowerSoil Pro HT protocol

The PowerSoil Pro HT were selected for further improvement based on the consistent DNA extraction quality and community profile for all soil types and its semi-automated QIAcube solution. Specifically, the effect of sample input amount as well as bead-beating intensity and time on the DNA quality, quantity, and community profile were tested on three different soil types in triplicates. Samples were processed as standard amplicon library preparations.

##### Sample input amount

Total DNA extracted was significantly greater for 125 mg sample input (ANOVA on ranks + Tukey's HSD: 16.4 % variance, $p<0.001$). Input amount did not affect the Shannon diversity index (ANOVA on ranks, $p=0.09$) or Bray Curtis dissimilarity (ANOVA on ranks, $p=0.62$) (Table 1).

Bray-Curtis dissimilarity of replicates within protocols was not different to the dissimilarity of replicates between protocols for any soil type.

Generally, the microbial community profiles were similar between sample input amounts (Fig 2A). Differential abundance analysis of ASVs with a relative abundance above 0.1 % revealed two ASVs to be differentially abundant in the Organic sample with 125 mg input (Fig 2D).

Principal Component Analysis (PCA) revealed the community profile clustered according to soil type, not input amount. Based on a PERMANOVA sample input amount did not explain any of the observed variance (Fig 3A). When stratifying for soil type PCA revealed samples clustered by sample input amount, however the explained variance was not significant in any case (Fig 3B-F).

**Table 1. General statistics from different sample input amounts.** All samples were rarefied to 3,511 reads (lowest read count in any sample with more than 3,000 total reads). ASVs not exceeding 0.1 % relative abundance in at least one sample were removed prior to Hellinger transformation and calculation of Bray-Curtis dissimilarity. Numbers represent mean and numbers in parentheses represent standard deviation.

| | | Laboratory Metrics (n=3) | | | Sequencing Metrics (n=Libraries) | | | | | | |
|---|---|---|---|---|---|---|---|---|---|---|---|
| **Soil type** | **Input [mg]** | **DNA yield [µg]** | **260/280** | **260/230** | **Libraries** | **Library conc. [ng/µL]** | **Number of reads** | **Observed ASVs** | **Shannon Diversity** | **Bray-Curtis Dissimilarity** | **BC Dissimilarity Between Input** |
| Beach Sand | 50 | 0.09 (0.03) | 1.09 (0.10) | 0.50 (0.09) | 3 | 5.01 (0.85) | 6492 (1589) | 2458 ( 5) | 7.64 (0.01) | 0.30 (0.02) | 0.34 (0.02) |
| | 125 | 0.16 (0.07) | 1.21 (0.07) | 0.55 (0.10) | 3 | 3.92 (1.20) | 4314 ( 834) | 2404 ( 51) | 7.60 (0.04) | 0.31 (0.01) | |
| Clay | 50 | 8.49 (1.65) | 1.88 (0.02) | 2.01 (0.03) | 3 | 14.44 (2.09) | 18970 (2452) | 2581 ( 14) | 7.67 (0.01) | 0.29 (0.01) | 0.29 (0.01) |
| | 125 | 17.63 (2.64) | 1.90 (0.00) | 2.12 (0.02) | 3 | 8.84 (7.48) | 7414 (5359) | 2563 ( 48) | 7.67 (0.04) | 0.30 (0.01) | |
| Organic | 50 | 7.66 (2.08) | 1.90 (0.02) | 2.00 (0.04) | 3 | 10.03 (1.87) | 20163 (3281) | 2907 ( 24) | 7.89 (0.01) | 0.48 (0.03) | 0.50 (0.03) |
| | 125 | 14.40 (4.18) | 1.83 (0.04) | 1.72 (0.32) | 3 | 14.69 (2.23) | 17742 (2182) | 2810 (105) | 7.84 (0.05) | 0.49 (0.08) | |
| Sand | 50 | 2.17 (0.68) | 1.77 (0.08) | 1.57 (0.26) | 3 | 9.63 (3.37) | 15542 (3477) | 2589 ( 18) | 7.73 (0.01) | 0.33 (0.02) | 0.35 (0.02) |
| | 125 | 6.32 (3.11) | 1.85 (0.03) | 1.87 (0.04) | 3 | 12.65 (1.11) | 18601 (2987) | 2566 ( 64) | 7.71 (0.03) | 0.34 (0.02) | |
| Sand-Clay | 50 | 2.97 (0.59) | 1.84 (0.04) | 1.74 (0.24) | 3 | 14.11 (0.53) | 19210 (2100) | 2541 ( 58) | 7.68 (0.04) | 0.35 (0.03) | 0.33 (0.03) |
| | 125 | 7.56 (0.89) | 1.83 (0.01) | 1.60 (0.22) | 3 | 10.52 (1.56) | 18993 (2423) | 2486 ( 29) | 7.64 (0.02) | 0.30 (0.01) | |

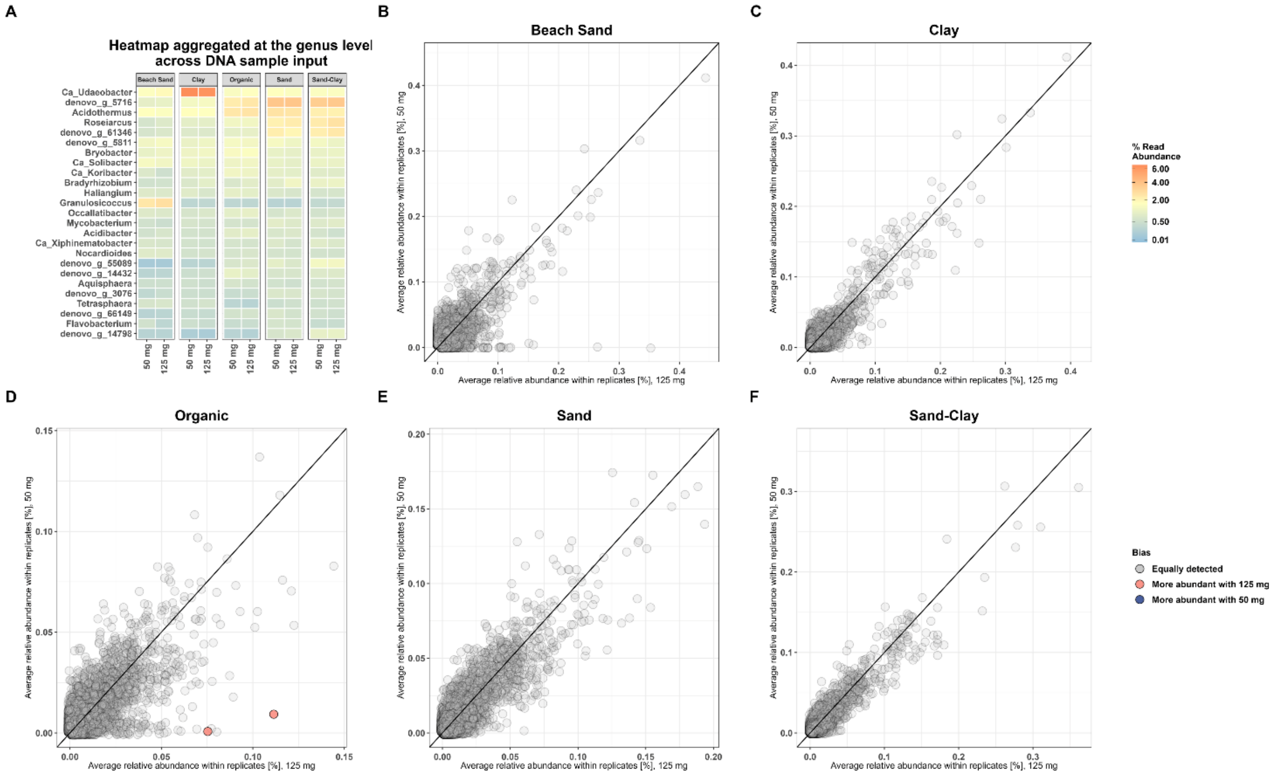


**Fig 2. Community characteristics and differential abundance plots based on amplicon data.** (A) Heatmap of community profile at genus level across sample input, faceted by soil type. Differential abundance plot for each soil type: (B) Beach Sand, (C) Clay, (D) Organic, (E) Sand, (F) Sand-Clay. ASVs differentially abundant in one protocol are marked with color. Bias was calculated with DESeq2 and was defined as a significant difference in log2-fold-change (adjusted p-value<0.05). ASVs filtered for minimum relative abundance >0.1 % in any sample.

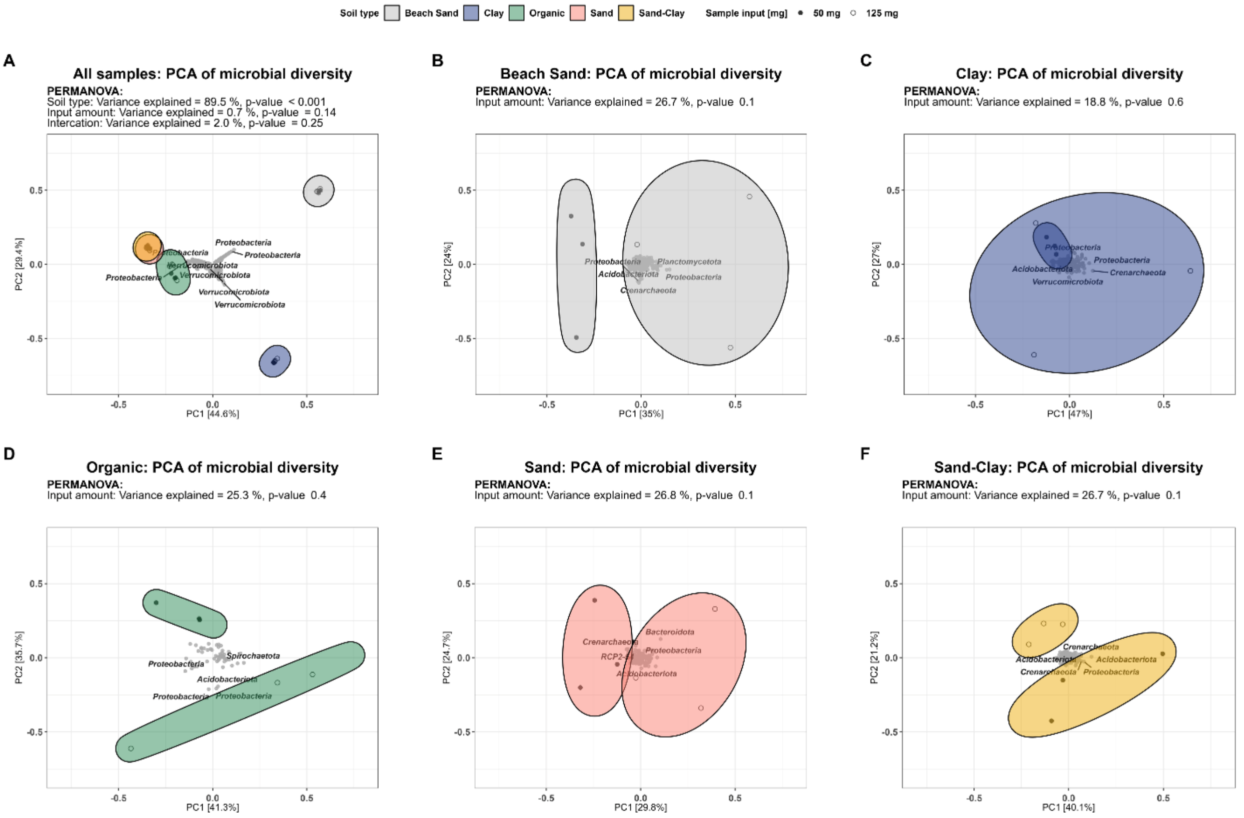


**Fig 3. PCA of DNA extraction with 50 mg input and 125 mg using PowerSoil Pro HT.** (A) PCA of DNA extraction with 50 mg input and 125 mg using PowerSoil Pro HT for all samples. Stratified by soil type: (B) Beach Sand, (C) Clay, (D) Organic, (E) Sand, (F) Sand-Clay. ASVs not exceeding 0.1% relative abundance in at least one sample were removed before Hellinger-transformation.

#### Bead beating intensity and time

The effect of bead-beating intensity and time on the DNA integrity and community characteristics were also tested for three soil types: Clay, Sand, and Organic. Specifically, four different bead-beating intensities 1200, 1400, 1600, and 1800 RPM were tested for three different bead-beating time durations: 2, 4, and 6 minutes. General stats can be found in table 2.

Both bead beating time and intensity had a positive correlation with the total DNA extracted, however no significant difference was observed between 1600 and 1800 RPM. Intensity explained 31.9% of variance (ANOVA on ranks, $p<0.001$, $n=108$), whereas time and soil type explained 14.0% ($p<0.001$) and 27% ($p<0.001$) respectively. All interaction terms combined explained 19.4 % of the variance ($p<0.001$). Generally, the 260/280 and 260/230 ratios increased with both time and intensity, most likely due to the increase in yield.

Peak fragment length was negatively correlated with both intensity and time. Intensity explained 54.9 % of variance ($p<0.001$), time explained 11.1 % of variance ($p<0.001$) and soil type explained 10.0 % of variance (p<0.001). The only significant interactions were between intensity and time (Explained variance: 4.3 %, p<0.001) and intensity and soil type (Explained variance: 5.2 %, p<0.001) (Table 2).

Bead beating intensity and time had little effect on the Shannon diversity index, except for Sand, for which the index stabilized at 1600 RPM. The microbial community of Sand and Clay had lower Shannon diversity index at low bead beating intensities and time (Table 2). Data for sand at 1200 RPM and 2 min is absent due to low read count as a result of poor amounts of DNA template. Bead beating intensity and time had little effect on the community profile of Organic soil (Fig 4A).

In PCA, samples are clustered according to soil type (Fig 4B). Stratified PERMANOVA constrained by time and intensity revealed that an average of 9.3 % of variance could be explained by intensity, and an average of 15.1 % of the variation could be explained by time (Fig 4C-E). For Organic soil, the RDA1 only accounted for 5.2 % of the total variation, thus the observed microbial community was less affected by bead beating time and intensity compared to Clay (10.5 %) and Sand (14.9 %). Furthermore, at or above 1600 RPM samples clustered regardless of bead beating time for Clay and Sand.

No ASVs were found to be differentially expressed when comparing 1600 RPM for 4 and 6 minutes of 1800 RPM for 4 and 6 minutes (Fig 5). Based on these results bead beating should be conducted at 1600 RPM and a total time of six minutes - two minute bead beating intervals separated by two minutes of incubation on ice.

**Table 2. General statistics from different bead beating protocols.** All samples were rarefied to 3,118 reads (lowest read count in any sample with more than 3,000 total reads). ASVs not exceeding 0.1 % relative abundance in at least one sample were removed prior to Hellinger transformation and calculation of Bray-Curtis dissimilarity. Numbers represent mean and numbers in parentheses represent standard deviation.

| | | | Laboratory Metrics (n=3) | | | | Sequencing Metrics (n=Libraries) | | | | | |
|---|---|---|---|---|---|---|---|---|---|---|---|---|
| **Soil type** | **Intensity [RPM]** | **Time [min.]** | **DNA yield [µg]** | **260/280** | **260/230** | **Integrity [kbp]** | **Libraries** | **Library conc. [ng/µL]** | **Number of reads** | **Observed ASVs** | **Shannon Diversity** | **Bray-Curtis Dissimilarity** |
| Clay | 1200 | 2 | 0.02 (0.01) | 1.57 (0.09) | 0.42 (0.29) | NaN ( NA) | 3 | 4.56 ( 1.91) | 5570 ( 303) | 2082 ( 48) | 7.42 (0.04) | 0.42 (0.01) |
| | | 4 | 0.19 (0.21) | 1.78 (0.15) | 0.90 (0.31) | 15.18 (2.43) | 3 | 26.11 (15.81) | 5346 ( 228) | 2335 (143) | 7.61 (0.10) | 0.44 (0.02) |
| | | 6 | 3.03 (0.07) | 1.90 (0.02) | 1.97 (0.07) | 12.17 (0.49) | 3 | 27.17 ( 4.85) | 4870 ( 971) | 2348 ( 14) | 7.61 (0.01) | 0.40 (0.00) |
| | 1400 | 2 | 0.38 (0.61) | 1.73 (0.08) | 0.79 (0.28) | 24.15 ( NA) | 3 | 18.34 (24.18) | 5256 (1131) | 2187 (219) | 7.50 (0.16) | 0.45 (0.05) |
| | | 4 | 0.78 (1.13) | 1.69 (0.30) | 1.18 (0.66) | 13.75 (1.08) | 3 | 26.84 (10.80) | 6473 ( 845) | 2341 ( 37) | 7.61 (0.03) | 0.41 (0.01) |
| | | 6 | 6.06 (1.73) | 1.91 (0.04) | 1.96 (0.11) | 10.88 (0.66) | 3 | 19.97 ( 9.58) | 5092 ( 571) | 2359 ( 17) | 7.61 (0.01) | 0.39 (0.02) |
| | 1600 | 2 | 2.93 (0.18) | 1.90 (0.01) | 1.96 (0.06) | 11.94 (1.49) | 3 | 30.55 ( 6.47) | 6171 ( 830) | 2338 ( 8) | 7.60 (0.00) | 0.38 (0.01) |
| | | 4 | 4.28 (0.16) | 1.89 (0.00) | 2.02 (0.09) | 12.89 (1.83) | 3 | 27.89 ( 3.38) | 6227 ( 952) | 2304 ( 29) | 7.56 (0.02) | 0.36 (0.01) |
| | | 6 | 5.54 (0.78) | 1.92 (0.01) | 2.12 (0.03) | 10.15 (0.65) | 3 | 24.14 ( 1.80) | 5865 ( 857) | 2354 ( 11) | 7.60 (0.00) | 0.38 (0.00) |
| | 1800 | 2 | 3.16 (0.22) | 1.89 (0.02) | 2.01 (0.04) | 11.05 (1.45) | 3 | 25.38 ( 7.82) | 6756 ( 926) | 2346 ( 23) | 7.60 (0.01) | 0.38 (0.01) |
| | | 4 | 5.97 (0.79) | 1.91 (0.01) | 2.11 (0.02) | 9.33 (1.98) | 3 | 27.43 ( 2.93) | 6308 ( 561) | 2321 ( 18) | 7.57 (0.01) | 0.37 (0.02) |
| | | 6 | 5.28 (0.35) | 1.90 (0.01) | 2.06 (0.02) | 9.28 (0.69) | 3 | 24.94 ( 4.64) | 6205 ( 497) | 2370 ( 14) | 7.61 (0.01) | 0.39 (0.01) |
| Organic | 1200 | 2 | 0.58 (0.16) | 1.82 (0.08) | 1.12 (0.48) | 14.74 (0.52) | 3 | 25.70 (20.13) | 7085 (1469) | 2642 ( 19) | 7.80 (0.01) | 0.54 (0.01) |
| | | 4 | 0.22 (0.05) | 1.71 (0.16) | 1.02 (0.09) | 12.64 ( NA) | 3 | 33.34 ( 9.84) | 6873 ( 589) | 2663 ( 8) | 7.82 (0.00) | 0.53 (0.02) |
| | | 6 | 0.51 (0.05) | 1.75 (0.01) | 1.29 (0.09) | 11.20 (1.75) | 3 | 42.58 ( 2.00) | 6821 (1151) | 2680 ( 4) | 7.83 (0.00) | 0.53 (0.02) |
| | 1400 | 2 | 1.14 (0.36) | 1.89 (0.09) | 1.20 (0.97) | 12.59 (0.57) | 3 | 27.94 (14.27) | 5988 ( 670) | 2638 ( 12) | 7.80 (0.00) | 0.50 (0.00) |
| | | 4 | 1.93 (0.51) | 1.92 (0.06) | 1.91 (0.16) | 12.81 (1.36) | 3 | 41.65 ( 8.48) | 5552 (1327) | 2687 ( 6) | 7.83 (0.01) | 0.52 (0.01) |
| | | 6 | 1.55 (0.21) | 1.85 (0.01) | 1.59 (0.27) | 9.52 (0.39) | 3 | 32.61 ( 2.44) | 5968 ( 834) | 2704 ( 10) | 7.84 (0.01) | 0.53 (0.02) |
| | 1600 | 2 | 1.19 (0.99) | 1.89 (0.04) | 1.88 (0.02) | 10.18 (0.60) | 3 | 36.62 ( 4.44) | 5703 ( 166) | 2671 ( 17) | 7.82 (0.01) | 0.53 (0.02) |
| | | 4 | 1.94 (0.12) | 1.85 (0.01) | 1.82 (0.09) | 8.56 (0.69) | 3 | 25.09 ( 2.27) | 7863 (1290) | 2667 ( 2) | 7.82 (0.00) | 0.53 (0.02) |
| | | 6 | 2.48 (0.19) | 1.85 (0.02) | 1.75 (0.24) | 9.77 (0.47) | 3 | 22.31 ( 2.04) | 6332 (2731) | 2668 ( 35) | 7.82 (0.02) | 0.50 (0.01) |
| | 1800 | 2 | 0.92 (0.02) | 1.89 (0.06) | 1.79 (0.12) | 9.11 (1.30) | 3 | 27.67 ( 2.41) | 5241 (2197) | 2654 ( 17) | 7.81 (0.01) | 0.50 (0.02) |
| | | 4 | 1.93 (0.26) | 1.91 (0.04) | 1.58 (0.46) | 8.00 (0.43) | 3 | 26.56 ( 2.39) | 5584 (1886) | 2690 ( 18) | 7.83 (0.01) | 0.52 (0.03) |
| | | 6 | 1.75 (1.12) | 1.84 (0.13) | 1.64 (0.31) | 7.05 (0.34) | 3 | 11.69 (10.14) | 6991 (1555) | 2662 ( 22) | 7.82 (0.02) | 0.54 (0.01) |
| Sand | 1200 | 2 | 0.02 (0.00) | 1.71 (0.23) | 0.56 (0.02) | NaN ( NA) | 0 | 0.52 ( 0.37) | NaN ( NA) | NaN ( NA) | NaN ( NA) | NA (NA) |

| | | | | | | | | | | | |
|---|---|---|---|---|---|---|---|---|---|---|---|
| | 4 | 0.02 (0.00) | 1.57 (0.16) | 0.63 (0.04) | NaN ( NA) | 3 | 3.43 ( 1.26) | 5814 ( 588) | 2015 (128) | 7.38 (0.14) | 0.43 (0.03) |
| | 6 | 0.41 (0.13) | 1.62 (0.04) | 1.03 (0.16) | 14.04 (2.59) | 3 | 42.20 ( 0.42) | 5408 ( 954) | 2420 ( 40) | 7.67 (0.03) | 0.41 (0.01) |
| 1400 | 2 | 0.02 (0.00) | 1.41 (0.08) | 0.59 (0.02) | NaN ( NA) | 3 | 2.68 ( 0.79) | 6045 ( 281) | 2012 (168) | 7.35 (0.13) | 0.48 (0.03) |
| | 4 | 0.03 (0.01) | 1.50 (0.03) | 0.64 (0.05) | NaN ( NA) | 2 | 6.21 ( 2.03) | 4892 ( 658) | 2114 ( 42) | 7.47 (0.03) | 0.44 (NA) |
| | 6 | 0.49 (0.28) | 1.82 (0.22) | 0.90 (0.76) | 12.48 (0.57) | 3 | 31.18 ( 0.85) | 6766 (2076) | 2381 ( 30) | 7.65 (0.02) | 0.41 (0.02) |
| 1600 | 2 | 0.50 (0.03) | 1.71 (0.06) | 1.22 (0.22) | 12.80 (1.74) | 3 | 28.68 ( 6.01) | 6420 (1862) | 2352 ( 12) | 7.62 (0.01) | 0.40 (0.02) |
| | 4 | 0.80 (0.04) | 1.78 (0.01) | 1.47 (0.13) | 13.81 (0.68) | 3 | 29.69 ( 2.86) | 6559 ( 685) | 2357 ( 13) | 7.63 (0.01) | 0.37 (0.01) |
| | 6 | 1.32 (0.31) | 1.85 (0.06) | 1.77 (0.06) | 12.32 (1.41) | 3 | 26.03 ( 5.34) | 5593 (2443) | 2361 ( 50) | 7.64 (0.03) | 0.39 (0.01) |
| 1800 | 2 | 0.49 (0.03) | 1.78 (0.05) | 1.43 (0.15) | 7.94 (0.30) | 3 | 27.00 ( 3.49) | 8170 (1670) | 2361 ( 28) | 7.64 (0.02) | 0.37 (0.02) |
| | 4 | 0.97 (0.17) | 1.89 (0.09) | 1.63 (0.12) | 8.81 (2.49) | 3 | 24.23 ( 2.26) | 7334 (1202) | 2371 ( 12) | 7.65 (0.01) | 0.40 (0.01) |
| | 6 | 1.40 (0.13) | 1.85 (0.07) | 1.86 (0.10) | 8.21 (1.14) | 3 | 26.18 ( 4.68) | 6622 (1251) | 2350 ( 4) | 7.63 (0.00) | 0.40 (0.01) |

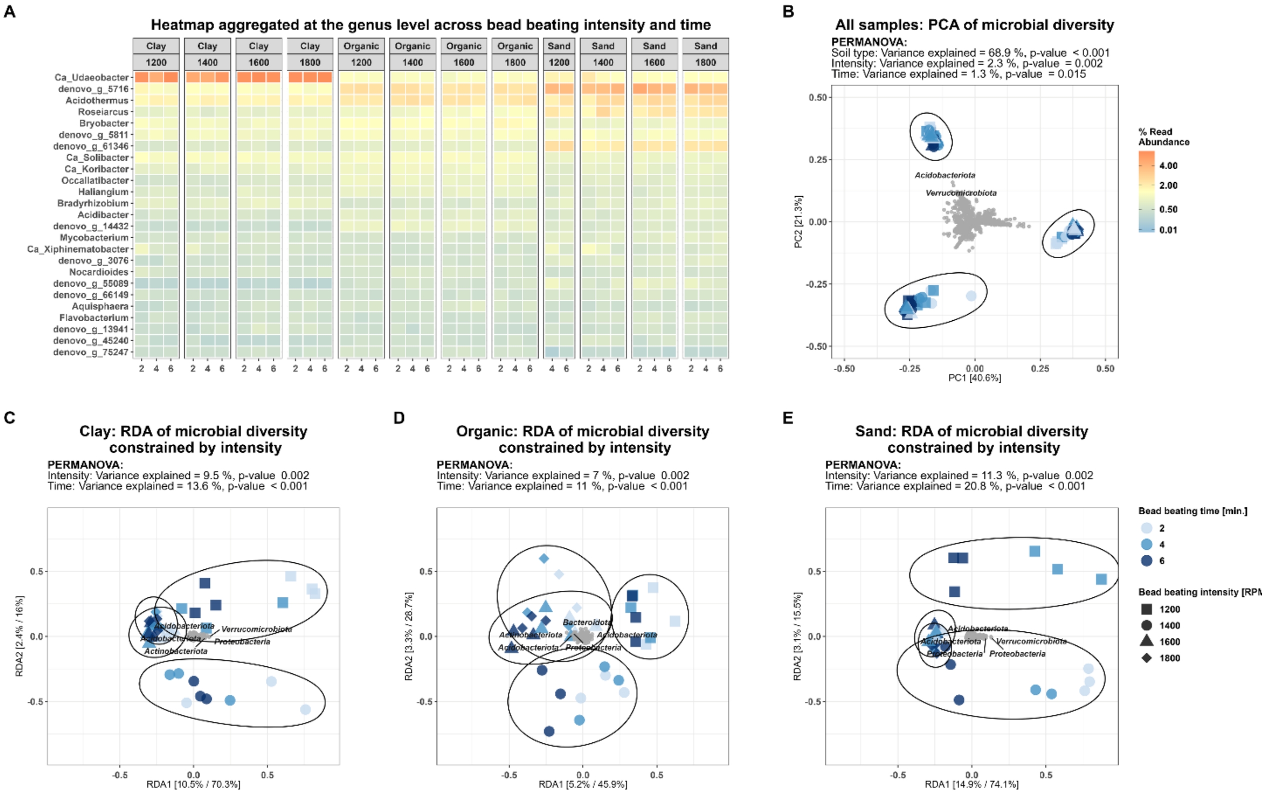


**Fig 4. Effect of bead beating intensity and total time on the community profile**. Community profile characteristics for the soil types, organic, clay, and sand. (A) Heatmap of community profile at phylum level across bead-beating time, faceted by soil type and intensity. (B) PCA of soil types at different bead-beating intensities and duration. RDA constrained by time and bead beating intensity stratified by soil types: (C) Clay, (D) Organic, and (E) Sand.

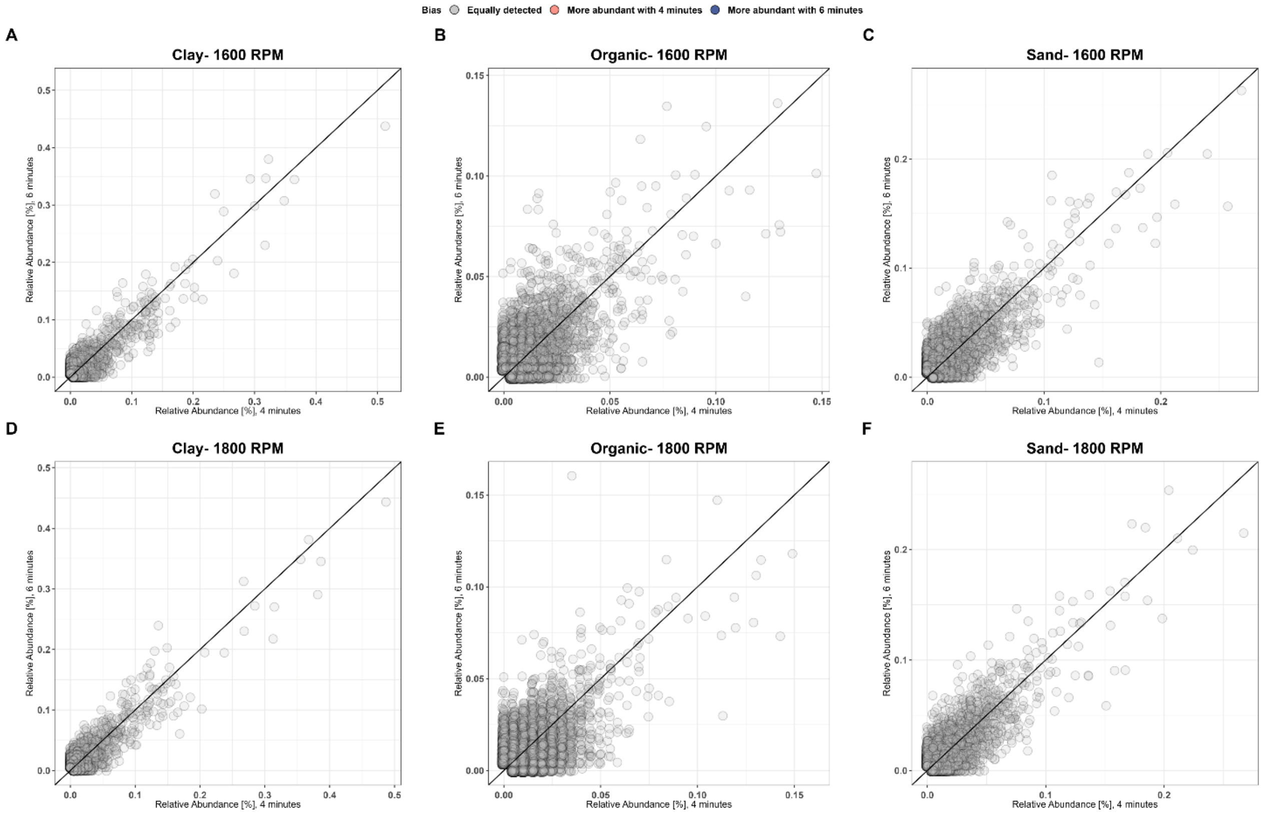


**Fig 5. Differential abundance plots at different RPMs.** Differential abundance plots with 1600 RPM for 4 and 6 minutes of bead-beating time stratified by soil type: (A) Clay, (B) Organic, (C) Sand. Differential abundance plots with 1800 RPM for 4 and 6 minutes of bead-beating time stratified by soil type: (D) Clay, (E) Organic, (F) Sand. Bias was calculated with DESeq2 and is defined as a significant difference in log2-fold-change (adjusted p-value<0.05). ASVs filtered for minimum relative abundance >0.1 % in any sample.
