## Supplementary material for "High-throughput DNA extraction and cost-effective miniaturized metagenome and amplicon library preparation of soil samples for DNA sequencing": S3 file

### Supplementary File 3:

#### Optimization of Illumina DNA prep

The Illumina DNA prep Reference Guide states the diluted metagenomic libraries should be size-selected with 45 µL SPB (IPB in the current kit) during the first step and 15 µL the second step. A successful two-sided clean-up should yield a peak fragment length of around 600 bp, however during the development of the miniaturized protocol, a shift in peak fragment length towards 400-500 bp was observed. Similar results have been reported by the application note for Illumina Nextera XT library prep using the Echo 525 Liquid Handler (1). Considering the bell-shaped fragment distribution and the combined length of the i7 and i5 IDT for Illumina Nextera DNA UD Indexes is 135 bp, a large proportion of the reads will be shorter than 265-365 bp. This is suboptimal for the sequencing of 2 x 150 bp read length of the NovaSeq 6000 platform.

Reducing the size selection volume with a factor of 10 would require pipetting 1.5 µL of SPB in the second size selection step, which was not feasible due to the viscosity. Hence, the PCR reactions were diluted with NFW to increase the SPB volume for easier handling. Adding 12 µL NFW to the 5 µL PCR product with 9 µL SPB in step 1 and 3 µL SPB in step 2 resulted in a peak fragment length of 429 bp (n=3) (Table 1/Fig 1, alias 9.0/3.0). Reducing the volume of SPB to 8 µL in the first step (Table 1, alias 8/3.0) resulted in a mean peak fragment length of 570 bp (n=3) while retaining sufficient library material. To make pipetting more consistent, the SPB volumes were doubled to 16 µL and 6 µL, and the NFW volume was correspondingly increased to keep the ratios constant. This resulted in a similar peak fragment length of 599 bp (Table 1, alias 16/6.0).

**Table 1. Experiment characteristics and results.** The size-selection volumes of PCR product, NFW, supernatant, and SPB mixed for step 1 and 2. The alias refers to the result displayed in Fig 1.

| | Step 1 | | | Step 2 | | Result | |
|---|---|---|---|---|---|---|---|
| Alias | PCR | NFW | SPB | Supernatant | SPB | Concentration [ng/µL] | Peak [bp] |
| Standard | 45 | 40 | 45 | 125 | 15 | - | - |
| 9/3.0 | 5 | 12 | 9 | 25 | 3 | 2.08 | 429 |
| 9/2.5 | 5 | 12 | 9 | 25 | 2.5 | 1.06 | 466 |
| 9/2.0 | 5 | 12 | 9 | 25 | 2 | 1.17 | 503 |
| 8/3.0 | 5 | 12 | 8 | 25 | 3 | 1.16 | 570 |
| 16/6.0 | 5 | 35 | 16 | 50 | 6 | 1.21 | 599 |

| | | | | | | | |
|---|---|---|---|---|---|---|---|
| 8/2.5 | 5 | 12 | 8 | 25 | 2.5 | 0.84 | 646 |
| 8/2.0 | 5 | 12 | 8 | 25 | 2 | 0.52 | 775 |
| 7/3.0 | 5 | 12 | 7 | 25 | 3 | 1.05 | 452 |
| 7/2.5 | 5 | 12 | 7 | 25 | 2.5 | 0.39 | 795 |
| 7/2.0 | 5 | 12 | 7 | 25 | 2 | 0.16 | 1204 |

The experiment was conducted on an Activated Sludge sample.

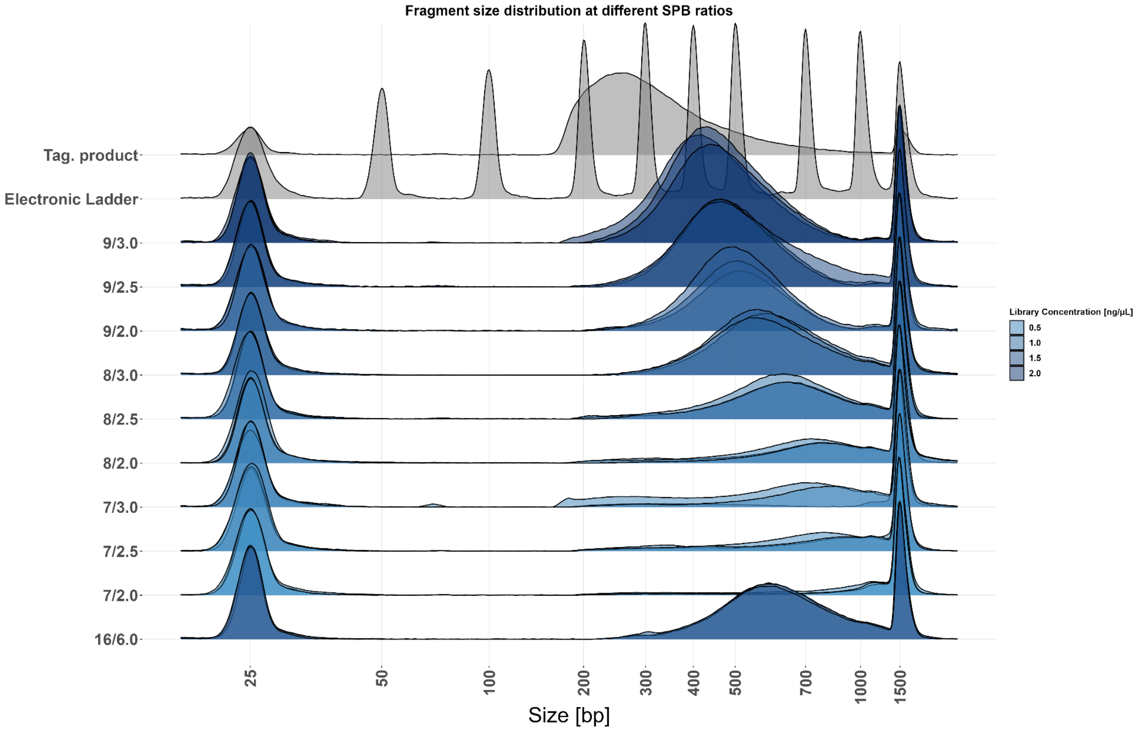


**Fig 1. Optimization of SPB bead ratios used during clean-up of metagenomic libraries**. Fragment size distribution of three individual replicates. Colored by average library concentrations in ng/µL from Qubit 1X HS DNA assays.

1. Effective Miniaturization of Illumina Nextera XT Library Prep for Microbiome Applications [Internet]. [cited 2023 Mar 23]. Available from: https://www.beckman.com/resources/reading-material/application-notes/effective-miniaturization-illumina-nextera-xt-library-prep-microbiome-applications
